# Quantitative measurement of phenotype dynamics during cancer drug resistance evolution using genetic barcoding

**DOI:** 10.1101/2025.03.26.645251

**Authors:** Frederick J.H. Whiting, Maximilian Mossner, Calum Gabbutt, Christopher Kimberley, Chris P Barnes, Ann-Marie Baker, Erika Yara, Andrea Sottoriva, Richard A. Nichols, Trevor A Graham

## Abstract

Cancer treatment frequently fails due to the evolution of drug-resistant cell phenotypes driven by genetic or non-genetic changes. The origin, timing, and rate of spread of these adaptations are critical for understanding drug resistance mechanisms but remain challenging to observe directly. We present a mathematical framework to infer drug resistance dynamics from genetic lineage tracing and population size data without direct measurement of resistance phenotypes. Simulation experiments demonstrate that the framework accurately recovers ground-truth evolutionary dynamics. Experimental evolution to 5-Fu chemotherapy in colorectal cancer cell lines SW620 and HCT116 validates the framework. In SW620 cells, a stable pre-existing resistant subpopulation was inferred, whereas in HCT116 cells, resistance emerged through phenotypic switching into a slow-growing resistant state with stochastic progression to full resistance. Functional assays, including scRNA-seq and scDNA-seq, validate these distinct evolutionary routes. This framework facilitates rapid characterisation of resistance mechanisms across diverse experimental settings.

## Introduction

The evolution of resistance remains the primary obstacle to the successful treatment of cancers. Initial studies focussed on resistance as a consequence of genetic alterations that produce resistant phenotypes that drive clonal evolution (Ding et al., 2012; Misale et al., 2012; Shi et al., 2014), but recent focus has shifted to non-genetic mechanisms of resistance (often termed ‘plasticity’) that enable cells to rapidly change phenotype (Emert et al., 2021; Hinohara et al., 2018; Rehman et al., 2021) promoting adaptive evolution following the change in selective pressure imposed by treatment. Efforts to tackle treatment resistance require knowledge of the molecular mechanism responsible so that it can be targeted. For example, new generations of targeted drugs use the resistance mechanisms of previous drugs as novel targets (Skoulidis et al., 2021). However, evolutionary-informed treatment strategies are also reliant on an understanding of the behaviour of a resistant phenotypes: approaches to “steer” tumour evolution to forestall or prevent resistance emergence require an understanding of the heritability of resistance mechanism(s) through cell divisions (Acar et al., 2020), and the relative fitness of resistant cells compared to drug sensitive counterparts (West et al., 2018).

Resistance evolution offers a unique opportunity to study phenotypic evolution in cancer: treatment is one of the few environmental changes a tumour experiences with known timing that is under clinical control, and resistance is clearly defined as a phenotype that survives treatment. These features mean studies into resistance are well placed to distinguish evolutionary behaviours such as the stability of a resistance phenotype through cell divisions and the environmental dependence of phenotypic change (Luria and Delbrück, 1943; Russo et al., 2022; Shaffer et al., 2020). These behaviours dictate how resistance is distributed amongst cell lineages over time which can in turn determine the relatedness of surviving cells.

In a patient tumour, naturally occurring somatic mutations can be interpreted as genetic barcodes (Gabbutt et al., 2022b). In an experimental setting, lineage tracing technologies enable the tracking of cell relatedness (Kebschull and Zador, 2018): unique genetic sequences are incorporated into cells’ genomes via lentivirus infection, meaning all subsequent ancestors of the parental, barcoded population’s cells inherit this experimentally measurable tag. Quantitative assessment of barcode data enables the measurement of the clonal dynamics (Blundell et al., 2019). When populations of cells are exposed to a change in extrinsic selection pressures, these genetic lineage tracing data provide a means to understand phenotype dynamics. For example, whether dominant lineages are shared between related experimental populations following exposure to drug treatment has been used as a qualitative indicator of whether drug resistance occurs via genetic or non-genetic mechanisms (Eyler et al., 2020).

However, there remain a lack of approaches that provide a quantitative readout of these phenotype behaviours. Here, we develop a mathematical modelling framework that infers the temporal dynamics of cancer cell drug resistance phenotypes using only genetic lineage tracing and population size data, without requiring specific measurement of cell phenotypes. We apply this framework to new experimental evolution data from two barcoded colorectal cancer cell lines that were exposed to long-term periodic chemotherapy, inferring distinct resistance phenotype dynamics between the two cell lines. We verify inferences made by the framework with additional functional interrogation of the cell lines including single-cell DNA and RNA sequencing.

## Results

### Quantitative models of resistance evolution

We developed mathematical models of resistance evolution that described the response of a population of cancer cells to periodic drug treatment. We aimed to infer the dynamics solely using the change in total population size and relatedness between surviving cell lineages. As such, we modelled the evolution of phenotypes within a cell population that control the response to treatment and the transitions between these phenotypes. Our approach was to begin with simple models of resistance evolution and add additional complexity only when simple models failed to explain data. We therefore designed three models of increasing complexity that encompass behaviours previously observed during the emergence of cancer cell resistance.

**Model A :** the first and simplest model (which we coin ‘unidirectional transitions’) consists of two phenotypes: sensitive and resistant. A ‘pre-existing resistance fraction’ parameter (*ρ*) controls the initial conditions of the model by setting the proportion of cells with the resistant phenotype when the model begins (*t_0_*). Cells can divide with phenotype-specific birth and death rates (sensitive: *b_S_,d_S_* and resistant: *b_R_,d_R_*). A ‘cost’ parameter (*δ*) controls a fitness penalty associated with the resistant phenotype in the untreated environment. Including a fitness penalty allows us to test the assumption that, in the absence of treatment, resistant cells exhibit a reduced net growth rate relative to sensitive cells: this behaviour has been observed in experimentally generated resistant cancer cell lines (Jensen et al., 2015), is significant for phenomena including the existence of slowly dividing ‘persister’ cells (Oren et al., 2021) and drug ‘addiction’ whereby resistant cells fail to grow in the absence of treatment (Maltas et al., 2024). Fitness penalties associated with a resistant phenotype also influences tumour containment strategies (Viossat and Noble, 2021). A switching parameter (*μ*) controls the probability of cells transitioning from the sensitive to resistant phenotype per cell division. Therapy was modelled by modifying the death rate of cells. For sensitive cells, the death rate increases as a function of *D(t)*, with *D(t)* representing the effective drug concentration at time t. Omitting the pharmacokinetics of the drug treatment would impair the model’s ability to recover observed changes in cell population sizes, as the cytotoxic effects of a drug are not realised immediately. We assume a pharmacokinetic model where the focus is on the change in the realized effect of the drug on cells, rather than directly modelling the drug concentration. The parameters *Dc* and *κ* regulate the strength and rate of accumulation/reduction of this effect, respectively. This model does not distinguish between intrinsic and extrinsic cellular factors; for instance, the increasing cytotoxicity experienced by cells following the addition of treatment could be affected by the drug’s chemistry, cellular efflux mechanisms, or some combination of the two. Furthermore, we make no assumption concerning the specific molecular change or mechanism that controls resistance in our models, and instead focus on the phenotypes as dictating the probability of survival during treatment. The parameter *ψ* models the survival probability of the resistant cells under treatment: *ψ = 1.0* denotes complete resistance, whereas when *ψ = 0.0* the sensitive and resistant cells experience the same level of drug-induced death.

To summarise, in this model the total cell population’s response to treatment is a product of the proportion of resistance (dictated by *ρ, δ* and *μ*), the effective strength of the drug at a given time (dictated by Dc and *κ*) and the strength of the resistant phenotype (dictated by *ψ*).

Model A allowed us to explore some simpler evolutionary scenarios of interest: compared to sensitive cells, resistant cells can either be absent or pre-exist at varying frequencies (by varying *ρ*), be either strongly or weakly resistant (by varying *ψ*) and can potentially grow at slower rates relative to sensitive cells in the untreated environment (by varying *δ*). Additionally, phenotypic switching from sensitive to resistant (*μ*) can vary, with lower probabilities representing the dynamics of genetic mutations and higher probabilities representing rapid non-genetic phenomena. Importantly, we did not define a strict boundary between these two possibilities, and instead can only reject genetic mutations when transitions occur at rates that are too high to be biologically feasible.

We considered which behaviours relevant to cancer resistance evolution model A (‘unidirectional switching’) could not reproduce. There are multiple reports of reversible transitions between resistant and sensitive phenotypes (Sharma et al., 2010), or the emergence of faster growing resistant cells from a slow cycling, drug tolerant population in response to the commencement of treatment (Russo et al., 2022). We therefore developed more complex models that could capture these dynamics.

Model B: ‘bidirectional transitions’ includes an additional behaviour whereby resistant cells can transition back to the sensitive phenotype with an independent probability (*σ*) per cell division. Previously, we had assumed resistance phenotypes arose unidirectionally (i.e. a resistant cell could not transition back to being sensitive). This modification allowed behaviours reminiscent of rapid, reversible, non-genetic transitions between phenotypes at higher probabilities, or forward and back mutations at lower probabilities.

Model C: ‘escape transitions’ - incorporates an additional phenotype we label ‘escape’. These cells behave like resistant cells in Model A and B but lack the fitness cost. As in model B, resistant cells can transition back to sensitive cells (with probability *σ*). However, resistant cells also now have a chance of transitioning to the escaped phenotype with a maximum probability of *α* per division. The transitions from resistant to escaped are scaled by a function of the current drug concentration (*α · f D(t)*) and are therefore only possible once treatment begins, permitting behaviours that resemble observations where transition rates and resistant phenotypes emerge in a drug-dependent manner (Pisco et al., 2013; Shaffer et al., 2017). Given transitions to the escape cells cannot commence until treatment begins, the initial conditions are still determined by the pre-existing resistance fraction parameter (*ρ*). With this modification the model can generate behaviours where the commencement of treatment allows slower growing cells (‘resistant cells’ in all three of our models) to transition to a faster growing phenotype also refractory to treatment (‘escape’ in model C).

Modelling Experimental Evolution: In all models, we coded-in features that enabled direct comparison to an empirical *in vitro* evolution experiment: the ability to track the lineage identity of cells uniquely labelled when the experiment commences (‘*in silico’* lineage tracing); a mutual expansion period before splitting cells into replicate sub-populations (where an initial barcoded pool of cell was split into replicate populations that were then independently evolved in parallel); exposing sub-populations to periodic drug treatment; and population sampling bottlenecks of known magnitude at given times. Our different models of resistance evolution - Models A,B,C – control the phenotypic composition of the cell populations during these sampling bottlenecks. The initial conditions (*ρ*) correspond to the composition of phenotypes when the simulations begin with N_0_ cells, when cells are also assigned a unique lineage identity (an *‘in silico’* barcode) (Fig.1). Cells from an expanded population (N_exp_) were sampled without replacement and allocated to drug-treatment replicates (DTx, where *x = 1, 2, 3, 4*). Additional sampling simulates the ‘passaging’ steps of evolution experiments (where P1 = Passage 1, etc.). Due to the limitation on population size imposed by the volume of tissue culture vessels *in vitro*, growth in all models was modelled as logistic growth with a fixed carrying capacity, C with a constant value guided by the empirical data. Cell populations were grown until a maximum population size (N_max_) was reached (where N_max_ *<* C). We also included the technical bottlenecks (e.g. sampling during barcode sequencing library generation) that occur in the laboratory setting and influence lineage tracing distributions (Blundell and Levy, 2014; Levy et al., 2015).

**Fig. 1.**
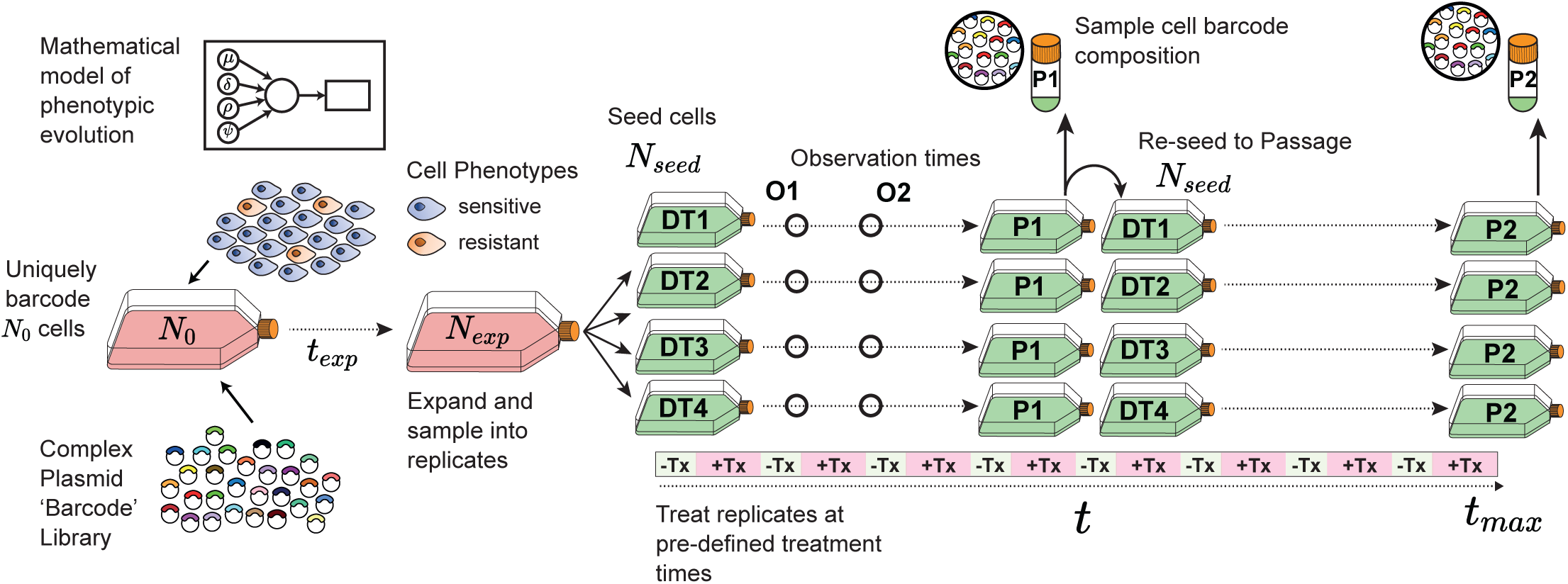
Overview of the experimental simulation and sampling procedure used to generate synthetic data. Unknown parameters govern the behaviour of cell phenotypes and their response to treatment. Fixed parameters control key experimental steps, including the initial number of uniquely barcoded cells, their expansion time, sampling bottlenecks when splitting into experimental replicates, and the timing of treatment windows. The drug-treatment replicates (DT1-4), population size observation timepoints (O1-2) and passages (P1-2) shown correspond to the experimental design used in our simulation modelling results.

We compare our models to existing mathematical models of phenotypic evolution in cancer. Model A shares features with commonly employed models of resistance evolution, incorporating two phenotypic compartments (sensitive and resistant), different pre-existing fractions of resistance, phenotype-specific birth and death rates (Diaz Jr et al., 2012; Foo and Michor, 2009; Misale et al., 2015) and modifications to these rates based on the current effective drug concentration (Chmielecki et al., 2011; Gallagher et al., 2024; Howard et al., 2018). Many studies allow for transitions from sensitive to resistant phenotypes, although this is often assumed to occur at a low probability, with values chosen based on estimates of mutation rates in cancer cells (Michor et al., 2005). While many models assume binary treated or untreated environments, fewer have directly incorporated drug pharmacokinetics to modify birth and death rates in a concentration-dependent manner (although see: Johnson et al., 2020; Poels et al., 2021).

Model B is comparable to previous theoretical work investigating how stochastic phenotypic switching can promote survival in bacteria (Acar et al., 2008; Tadrowski et al., 2018). These behaviours have also been modelled in cancer, where common features mirroring our own include asymmetrical transitions between phenotypes at probabilities too high to occur via genetic mutations (Dhawan et al., 2016; Nam et al., 2021). Importantly, we use our models to infer non-genetic phenotypic transitions by using these high probabilities to exclude genetic mutations. This ‘population-down’ approach contrasts with modelling strategies that predict non-genetic phenotypic transitions as a property of gene regulatory networks (Cheng et al., 2014; Hari et al., 2020).

Model C, our most complex model with three phenotypes, permits unidirectional transitions to a third phenotype, ‘escape,’ as a function of the current effective drug treatment (*α⋅fD(t)*), mirroring previous models that include drug-induced transitions from sensitivity to resistance (Kuosmanen et al., 2021; Pisco et al., 2013; Russo et al., 2022) and epigenetic rewiring to a stably resistant state that is drug-dependent and non-reversible (Gunnarsson et al., 2020).

In contrast to many existing models, we aim to infer multiple behaviours relevant to both drug treatment and resistance evolution from the data simultaneously, including the rate of change (*κ*), the strength of selection exerted by the drug (Dc), and the magnitude of resistance (*ψ*). Our modelling also explicitly accounts for the technical sampling steps encountered during long-term resistance evolution experiments which are particularly relevant in lineage tracing setups (Acar et al., 2020; Lan et al., 2017).

### Population and lineage distributions during treatment are determined by phenotype behaviours

We explored the population dynamics during treatment quantitatively using our modelling framework. We performed simulations with biologically feasible parameter values and recorded how replicate sub-populations responded to rounds of treatment. We hypothesised that phenotypic behaviours of interest - for example, transition rates between sensitive and resistant states and the proportion of resistance when treatment commences - could be inferred from empirical measurements. Each of our models was simulated to generate data that would typically be measured during a long-term evolution experiment. Per simulation, we modelled four drug-treatment replicates (DT1-4). These replicate cell populations had been sampled from the same expanded population or cells that had been uniquely labelled with *in silico* lineage tags - ‘barcodes’ - when the simulation began. Given a sufficient expansion period, closely related cells that share a lineage barcode should be present in multiple replicates. We also saved the total population size at two ‘observation’ timepoints (O1,O2) and each Passage step (P1,P2). We reasoned that comparing the change in total population size and the lineage identify of surviving subclones within and between replicates could be used to distinguish different evolutionary scenarios of interest. Initially we focussed on the change in the total number of cells over time.

We found that changes in total cell population sizes during treatment are primarily driven by the proportion of each phenotype and their response to treatment.

In model A, unsurprisingly, when resistance was initially more common (higher values of *ρ*), the time until the maximum population size was reached was shorter (Fig 2A vs 2B). Interestingly, even in cases where resistance was initially rare or absent, the changes in total cell population sizes between replicates can be highly repeatable (Fig 2B): despite the bottleneck exerted by treatment, the similar proportions of resistance in each replicate generated by transitions to resistance (*μ*) and the shared survival probability during treatment (*ψ*) generate consistent growth dynamics.

**Fig. 2.**
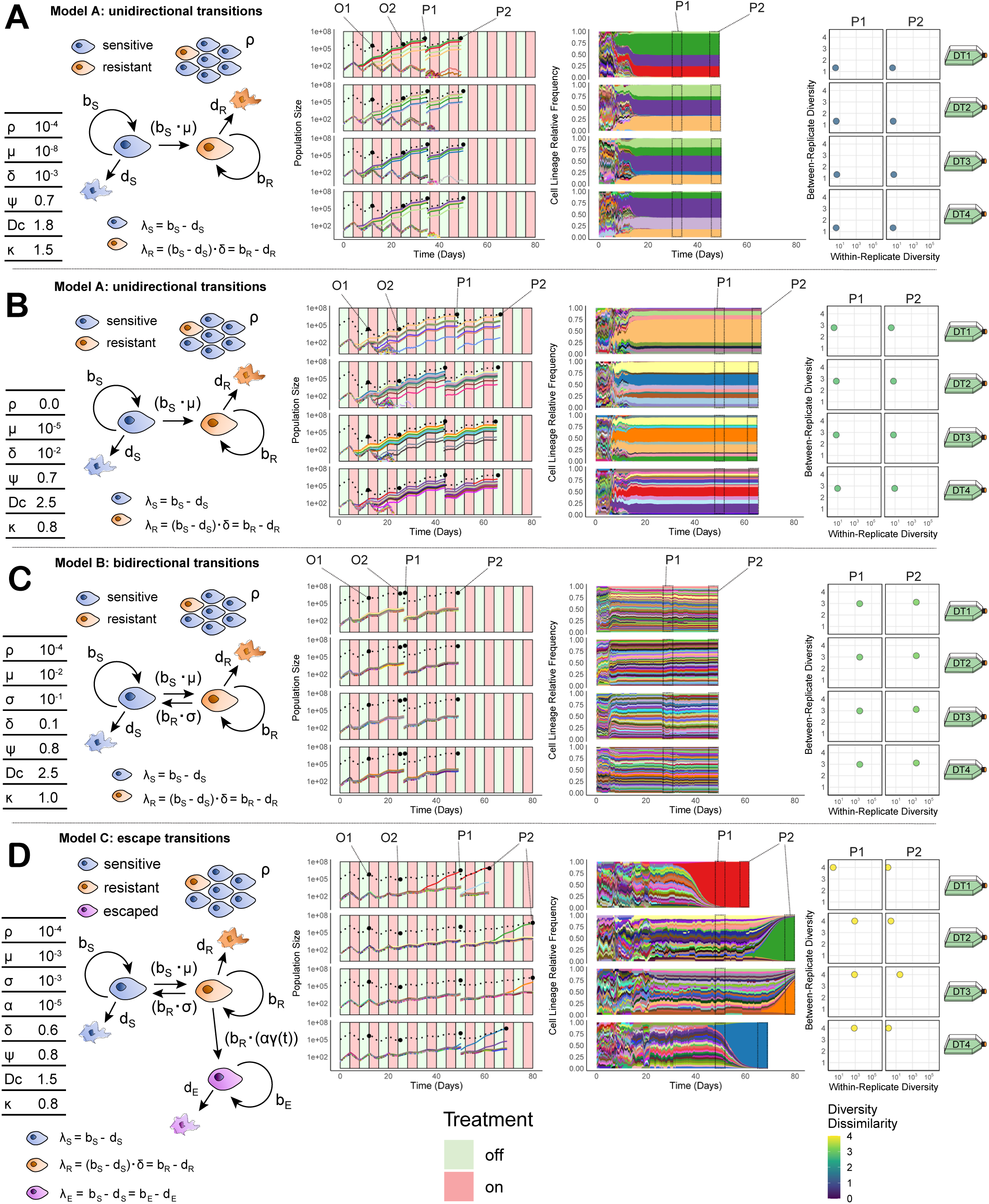
Quantitative models of resistance evolution exhibit distinct cell population dynamics during treatment. **A-D)** Per panel: (Left) Schematics of different evolutionary models, highlighting key parameters that control the behaviour of resistant, sensitive, and escape phenotypes (see text and Supplementary Theory Note for full descriptions). (Middle) Total cell population size trajectories (dashed lines) and the top 20 lineage population sizes (coloured lines)/relative frequencies (shaded areas) across four experimental drug-treatment replicates (DT1-4, rows). Lineage colours are consistent between replicates per panel. Four total cell size observations per replicate (two intermediate population size timepoints (O1 & O2) and two passage size timepoints (P1 & P2)) are highlighted. (Right) Lineage diversity statistics for the 4 drug-treatment replicates at two passage timepoints (P1 & P2 - columns), including within-replicate lineage (x-axis) and between-replicate diversity (y-axis). Simulation parameters: *b_x_* – birth rate of phenotype *x*, *d_X_* – death rate of phenotype *x*, *ρ* - pre-existing fraction of resistance, *μ* – sensitive to resistant transition probability per cell division, *σ* – resistant to sensitive transition probability per cell division, *α* – resistant to escape transition probability per cell division, *γ* – normalised effective drug concentration, *ψ* – strength of the resistant phenotype, Dc – maximum strength of the drug, *κ* – accumulation/decay rate of the drug.

In all models, if the effective phenotype transition probabilities were very low, the differing lag times preceding the emergence of treatment refractory populations could produce stochasticity in the time taken for the population to recover. This was especially true in model C (‘escape transitions’) when the transition from ‘resistant’ to ‘escape’ (*α*) was low: a population of slower growing resistant cells survive initial rounds of treatment before rarer transitions to the faster growing escape phenotype emerge during treatment (Fig 1D).

We next considered the signal in the lineage distributions (clone size distributions) over time. Phenotypic transitions alter the relationship between phenotype and the lineage identity assigned at the beginning of the experiment. When transitions are rare, lineage identity mirrors phenotype identity, but this correlation is progressively lost at higher phenotypic transition rates. Combined with the known sampling bottlenecks and other evolutionary parameters controlling the phenotype-specific growth rates and strength of selection exerted by treatment, these behaviours determine the identify of surviving lineages.

Current empirical lineage tracing systems enable tracking of over 10^6^ unique lineages. The sampling design we explored measures these lineage distributions 8 times per experiment (DT1-4 at P1-2). To enable the interpretable comparisons of these high-dimensional data, we condensed the lineage relationship distributions from each simulation into 2-dimensional diversity statistics that captured the diversity within replicates (x-axis) and the diversity between replicates (y-axis) simultaneously (Jasinska et al., 2020; Jost, 2006; Roswell et al., 2020) (Fig. 2 - right hand column).

Lineage tracing data gave extra information over-and-above population size data. For example, following treatment the repeated selection for the same lineages was indicative of stable, pre-existing resistance (Fig. 2A), whereas surviving lineages were more dissimilar between replicate sub-populations when rapid phenotypic transitions from sensitive to resistance occurred within separate lineages (Fig. 2B). Importantly, these differences in between-replicate diversity (y-axis) distinguish scenarios that led to similar levels of within-replicate resistance, and by extension within-replicate diversity (x-axis) (Fig. 2A & 2B - right hand columns), highlighting that information encoded in the lineage relationships offered additional means to reveal how resistance phenotypes were behaving. In cases where the rates of phenotypic switching were high, resistance was distributed across many lineages when treatment began, and the within-replicate diversity was also high (Fig. 2C). Stochasticity in the cell population size changes was often mirrored in the lineage diversity statistics: rapid expansions of ‘escape’ clones caused a crash in diversity where the emergence timing of the surviving lineage was unique to each replicate (Fig. 2D), increasing the variance in the corresponding within-replicate diversity axis (x axis). The identity of the lineage within which this escape transition occurred was also different between replicates, leading to high values of between-replicate diversity (Fig. 2D - right hand column).

In summary, the different evolutionary scenarios (encoded in different parameter values in our three models) lead to noticeable differences in the total cell population size and cell lineage changes throughout the treatment course.

### Population and lineage information can recover evolutionary parameters and different evolutionary scenarios

We next aimed to recover model parameters that describe the dynamics of resistance evolution using these population size and cell lineage relationship data. To achieve this, we developed an approximate Bayesian computation (ABC) inference approach.

We designed a hybrid phenotypic compartment approach to simulate the population size of each phenotype for each of our three models, treating the population as a stochastic jump process when below a predefined threshold and switching to a deterministic approximation when the population size reaches or exceeds this threshold. This approach preserves the important stochastic dynamics at small population sizes while efficiently simulating the system when populations are large (Germano et al., 2024; Kreger et al., 2021). We also constructed an agent-based model that preserves all the features of our first model. This second approach was more computationally expensive than our hybrid phenotypic compartment model, but also tracks the lineage identities of individual cells within the system. As such, this agent-based version could also generate cell lineage distributions.

Our fitting process consisted of two steps. First, we used the hybrid phenotypic compartment model to exclude parameter combinations that could not account for observed changes in cell population sizes during treatment by comparing vectors of population size changes at predetermined times (O1, O2, P1, P2). Next, we used our agent-based lineage model to simulate the remaining non-identifiable parameter combinations and generate cell lineage distributions at each passage (P1, P2). We approximated the posterior distribution by retaining parameter combinations that produced lineage distributions with diversity summary statistics (within- and between-replicate diversity) that closely matched the observed data. Having shown that they capture important features of the cell barcode distributions that are dictated by the underlying dynamics of resistance evolution, we used the within- and between-replicate lineage diversity statistics for this second step.

To assess the accuracy of our framework, we evaluated our ability to recover parameters from synthetic data where the true parameter values (ground truth) were known. For a specific set of parameters, we collected data on total population size changes and lineage distributions at predefined treatment times across four replicate sub-populations (Fig. 1).

When inferring parameter values without incorporating lineage distributions in the second step of our fitting procedure, we found that, while these observations could exclude many parameter combinations, a broad range of possibilities remained that the model could not distinguish (Supp. Fig. 1B). Whilst population trajectories alone allowed us to recover the effective strength of treatment (Dc), the parameters governing the distribution of the resistant phenotype within and between cell lineages over time (such as *μ* and ρ) were poorly identified. Specifically, when resistance was rare in the cell population, the initial drop in population size at the early observation timepoint (O1) was mainly driven by the selection pressure on sensitive cells (as determined by Dc). However, the subsequent population rebound during treatment, driven by the emergence of predominantly resistant cells, could be explained by different combinations of pre-existing resistance (*ρ*) and phenotypic transition probabilities from sensitive to resistant (*μ*).

When performing inference using both sources of information, the lineage relationships at each timepoint - as described by the diversity statistics - increased the power to identify evolutionary parameters markedly (Supp. Fig. 1D). For example, the addition of lineage information could distinguish between parameter combinations that led to similar overall proportions of resistance and sensitivity, but markedly different rates of phenotype switching (different *μ* values) and high fitness costs associated with resistance (different *δ* values) which cause selective lineage expansion and contraction (Fig. 2B-C). Repeating this model inference framework on parameter sets for each of our resistance models showed the approach could recover values from a range of evolutionary scenarios (Supp. Fig. 2).

There were scenarios that led to treatment having a minimal impact on the cell population size, limiting the modelling framework’s ability to recover the ground truth parameters. This was the case if resistance was extremely common (e.g. high values of *ρ*) or if the strength of selection exerted by the drug was weak (low values of Dc). Notably, whilst both behaviours lead to an unresponsive population, they do so due to subtly different reasons. If resistance is common, there can still be a small fraction of sensitive cells that are killed during treatment, whereas a weak drug effect would kill all cells with an equally low probability. Since our modelling framework relies on the population size changes and relationships between surviving lineages to identify likely parameter combinations, these scenarios hindered accurate parameter recovery (Supp. Fig. 3). These scenarios are less relevant to studying the dynamics of resistance evolution, as they represent situations where the population is essentially drug-resistant at the outset.

In model B, we found the framework usually failed to accurately identify the reversion switching probability from resistant to sensitive (*σ*) (Supp. Fig. 2B). Given this switching led to cells reverting to the sensitive phenotype within previously resistant cell lineages, they were often then lost in subsequent treatment rounds, obscuring the behaviour’s impact. In addition, higher values of the reversion switching probability (*σ*) have a similar impact on population composition as a lower strength of the resistant phenotype (*ψ*): cells that were previously resistant are killed independent of their lineage identity, either due to a rapid reversion to sensitivity or a weaker resistant phenotype. In more detail: phenotype switches back to a sensitive state (controlled by parameter *σ*) lead to an overall increased rate of cell death for the lineage where the switch has occurred, and a reduction in the size of the lineage. Because phenotype switches occur at a uniform rate across all clonal lineages, a higher rate of reversion back to the sensitive state therefore has an identical effect to increasing the direct death rate of the resistant cells (lower *ψ*). Incomplete resistance may occur due to very rapid phenotypic shifts, for example due to the stochastic partitioning of transcripts during cell division (Soltani et al., 2016).

The framework also struggled to recover the model behaviours if diversity was lost too rapidly. Our inference relies on distinguishing possible evolutionary scenarios by identifying shifts in diversity statistics between sampling steps. If the selection pressure exerted by treatment was too strong, only one/a few barcodes remained in each replicate by the first sampling step and the statistical power to distinguish scenarios was lost.

We explored whether these difficulties could be addressed with newer ‘evolving barcodes’ that continuously generate novel barcode tags within cell lineages, enabling higher resolution reconstruction of cell phylogenies (Kalhor et al., 2017). To explore the power of evolving barcodes in a simple setting we performed an additional re-barcoding step at the first passage (P1) to reintroduce measurable diversity into the system. We found that this information didn’t help solve the unidentifiability of the strength of resistance and reversion transition probability in Model B (*σ* vs *ψ*). However, in our Model C, it could help identify clonal sweeps due to ‘escape transitions’ that occurred following a complete loss of diversity prior to P1 (Supp. Fig. 4).

The difficulties in recovering the reversion phenotypic transition probability (*σ*) were absent for the sensitive to resistant transition (*μ*) where the probability strongly dictates the number and identity of surviving lineages. Given this unidentifiability of Model B (‘bidirectional transitions’), we chose to focus on the differences between models A and C - unidirectional switching & escape transitions.

As well as identifying parameters within each of our models, we wanted to know when the framework could distinguish between them. We therefore simulated two sets of ‘ground truth’ scenarios using each of our models A and C (28; 14 per model) and compared their ability to recover the true evolutionary scenario. Due to the differing complexity of our models (as captured by the differing number of parameters), we used the Deviance Information Criterion (DIC), a measure of model fit that penalises for model complexity (Francois and Laval, 2011).

We found there were cases where the differences left in the lineage distributions enabled the identification of the correct evolutionary model (as illustrated by a lower DIC score) (Supp. Fig. 5): the data generated by the simpler model A (‘unidirectional’) were correctly identified for all of the scenarios investigated, whilst in over half of the cases where the true model was model C (‘escape transitions’) the model A (‘unidirectional switching’) was preferred; the evidence in favour of our more complex model (model C) had to be strong to confidently reject our simpler model (model A). Rejection of model A occurred when the transition to the ‘escape’ phenotype in model C led to a stochastic crash in diversity that could not otherwise be explained by the total cell population size changes (Supp. Fig. 5B). These behaviours were observed when a moderate level of resistance (higher values of *μ* or *ρ*) with an associated fitness penalty (via *δ*) survived the initial rounds of treatment, and the probability of experiencing an escape transition (*α*) was relatively low.

In summary, for the scenarios explored here, we found that by combining commonly measured features of cell populations during long-term evolution experiments – including cell population size changes and cell lineage distributions (summarised via diversity statistics) - we could recover parameters that govern the response of the population to treatment and the evolution of resistance phenotypes. Furthermore, we show that in certain scenarios there are statistical signatures left by our more complex model that allow us to distinguish it from a simpler model characterised by two phenotypes and unidirectional switching.

### Experimental evolution of drug resistance in colorectal cancer

We applied our modelling framework to investigate the dynamics of resistance evolution in two common colorectal cancer cell line models: SW620 (a line exhibiting chromosomal instability (CIN)) and HCT116 (a mismatch repair deficient (MMRd) line).

To enable the tracking of cell lineage information, we labelled each cell line with a stable genetic barcode system (ClonTracer - Bhang et al., 2015). We refer to the barcoded cell lines as SW6bc (SW620) and HCTbc (HCT116). After growing cells for a short expansion step (approximately 6 population doublings in HCTbc and 5 population doublings in SW6bc) to ensure most barcodes were represented multiple times, we split each cell-line’s barcoded parental population (POT) into control (CO1-4) and drug treatment (DT1-4) replicates and subjected each subpopulation to periodic treatments of vehicle control (DMSO) or IC_50_ concentrations of 5-fluoruracil (5-Fu) (a constituent part of standard-of-care colorectal cancer chemotherapy treatments such as FOLFOX) (Fig. 4A & 5A). We isolated and sequenced barcoded single cell colonies to ensure the majority of cells contained a single barcode (Supp. Fig 6). We combined cell counts and imaging to derive total cell population size estimates at four time points (O1,O2,P1,P2) per treatment replicate (DT1-4) within the first two passages and extracted and sequenced the heritable genetic barcodes to measure the cell lineage distributions at each passage. We also used the distribution of lineages from the expanded, untreated cells (POT) to estimate the parental populations’ average birth and death rates for each cell line and used these estimates to parameterise the full resistance models. We then fit our models of resistance evolution to the total cell population size estimates and lineage distribution data. Because we observed sustained lag times between the commencement of treatment and the first passage (P1) in both cell lines (SW6bc: +37 days; HCTbc: +67 days - Fig. 4B & 5B) we were confident that resistance was not already the dominant phenotype and that the selective pressure exerted by treatment was significant.

**Fig. 3.**
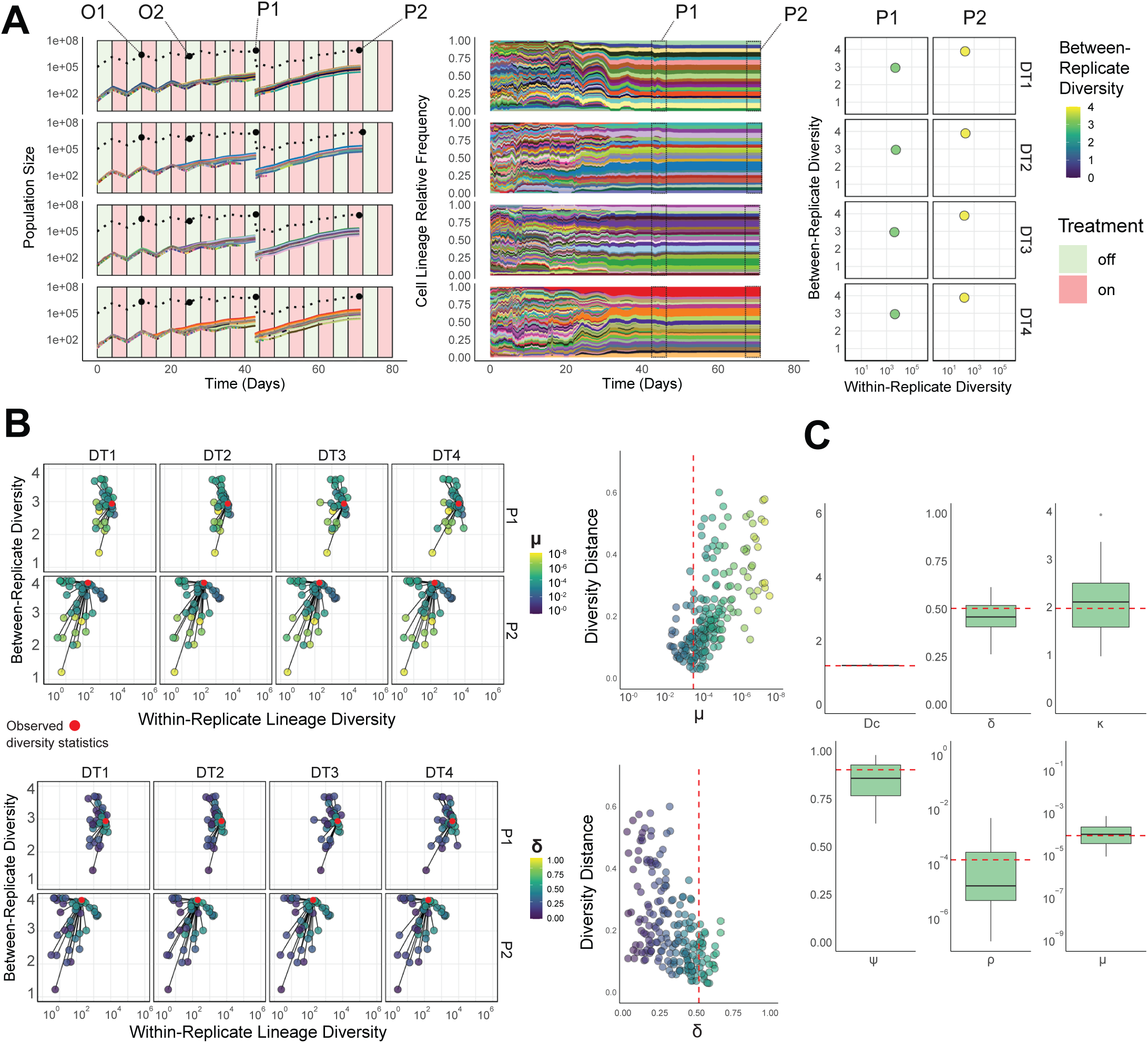
Lineage information enables the recovery of parameters controlling resistance evolution. **A)** Simulated population size (Left) and cell lineage (Middle) changes across 4 experimental drug-treatment replicates (DT1-4) and their lineage diversity statistics (Right). Total cell population size trajectories (dashed lines) and the top 20 lineage population sizes (coloured lines)/relative frequencies (shaded areas) across four experimental drug-treatment replicates (DT1-4, rows). Lineage colours are consistent between replicates. Four total cell size observations per replicate (two intermediate population size timepoints (O1 & O2) and two passage size timepoints (P1 & P2)) are highlighted. **B)** A subset of the simulated lineage diversity statistics for each sub-population’s two timepoints (P1-2, rows) used for the parameter inference step, highlighted by one of the model parameters (Top: sensitive to resistant phenotype transition probability per cell division - μ) and the fitness penalty of resistance (Bottom: controlled by δ) compared to the true diversity statistics for the given simulation (red points). Adjacent panels show the normalised lineage diversity distance as a function of the two parameters, with the true parameter value highlighted (red dashed line). **C)** Posterior distributions (n=50, 4 experimental replicates) of the inferred parameter values using the combined cell population size and cell lineage statistics (true parameter values - red dashed lines). Boxplots show the median, the first and third quartiles, and whiskers 1.5x the interquartile range.

**Fig. 4.**
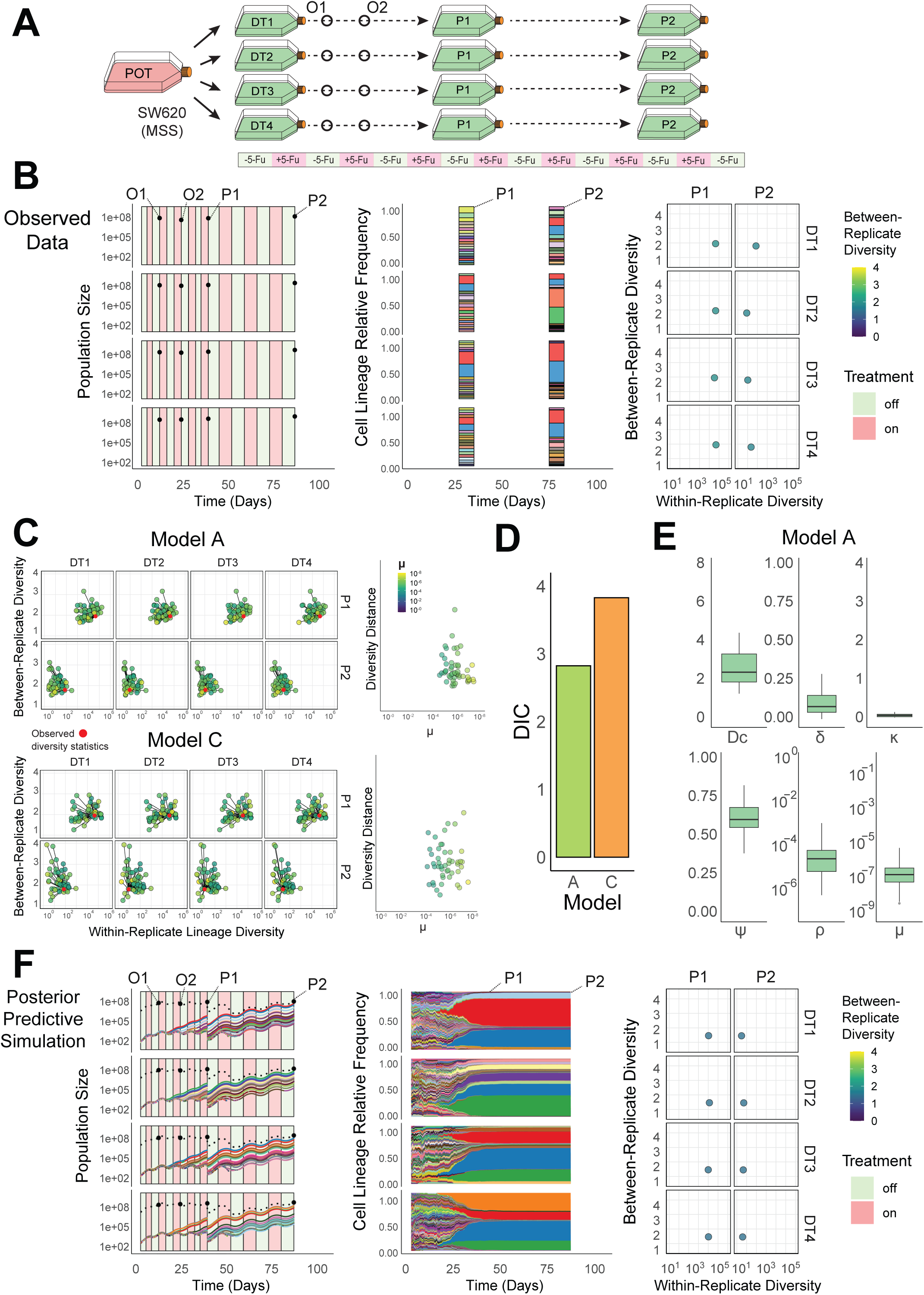
Quantification of the evolutionary dynamics during treatment in a barcoded colorectal cancer cell line (SW6bc). **A)** A simplified schematic of the experimental design used in a long-term evolutionary resistance assay. Barcoded SW620 cells (SW6bc) were barcoded and expanded (POT) before being sampled into four replicate drug-treatment sub-populations (DT1-4) that were exposed to periodic chemotherapy treatment (5-fluoruracil: 5- Fu) and passaged at two timepoints (P1 & P2). **B)** Population size counts at four timepoints per-replicate (DT1-4), including two intermediate population size estimates (O1 & O2) and two Passage timepoints (P1 & P2) (left panel); top 20 sequenced barcode lineage relative frequencies at the two passage timepoints, where area colour denotes lineage identity (middle panel); lineage diversity statistics of the sequenced barcode distributions at the two passage timepoints (P1 and P2) for each replicate composed of within replicate lineage diversity and between-replicate diversity dissimilarity. **C)** Posterior predictive simulations for Model A (unidirectional transitions – top) and Model C (escape transitions – bottom) of the within and between-replicate diversity statistics (LHS) and the normalised diversity distance from the observed statistics (red points) to the simulated values (highlighted by the transition parameter μ). **D)** The Deviance Information Criterion (DIC), a measure of model fit, for Model A & Model C (lower values indicate higher model support) calculated using the posterior predictive distributions. **E)** The posterior distributions (n=50, 4 experimental replicates) for parameters in Model A (the model with the highest support). Boxplots show the median, the first and third quartiles, and whiskers 1.5x the interquartile range. Parameters: *ρ* - pre-existing fraction of resistance, *μ* – sensitive to resistant transition probability per cell division, *ψ* – strength of the resistant phenotype, Dc – maximum strength of the drug, *κ* – accumulation/decay rate of the drug. **F)** A posterior predictive simulation using Model A.

**Fig. 5.**
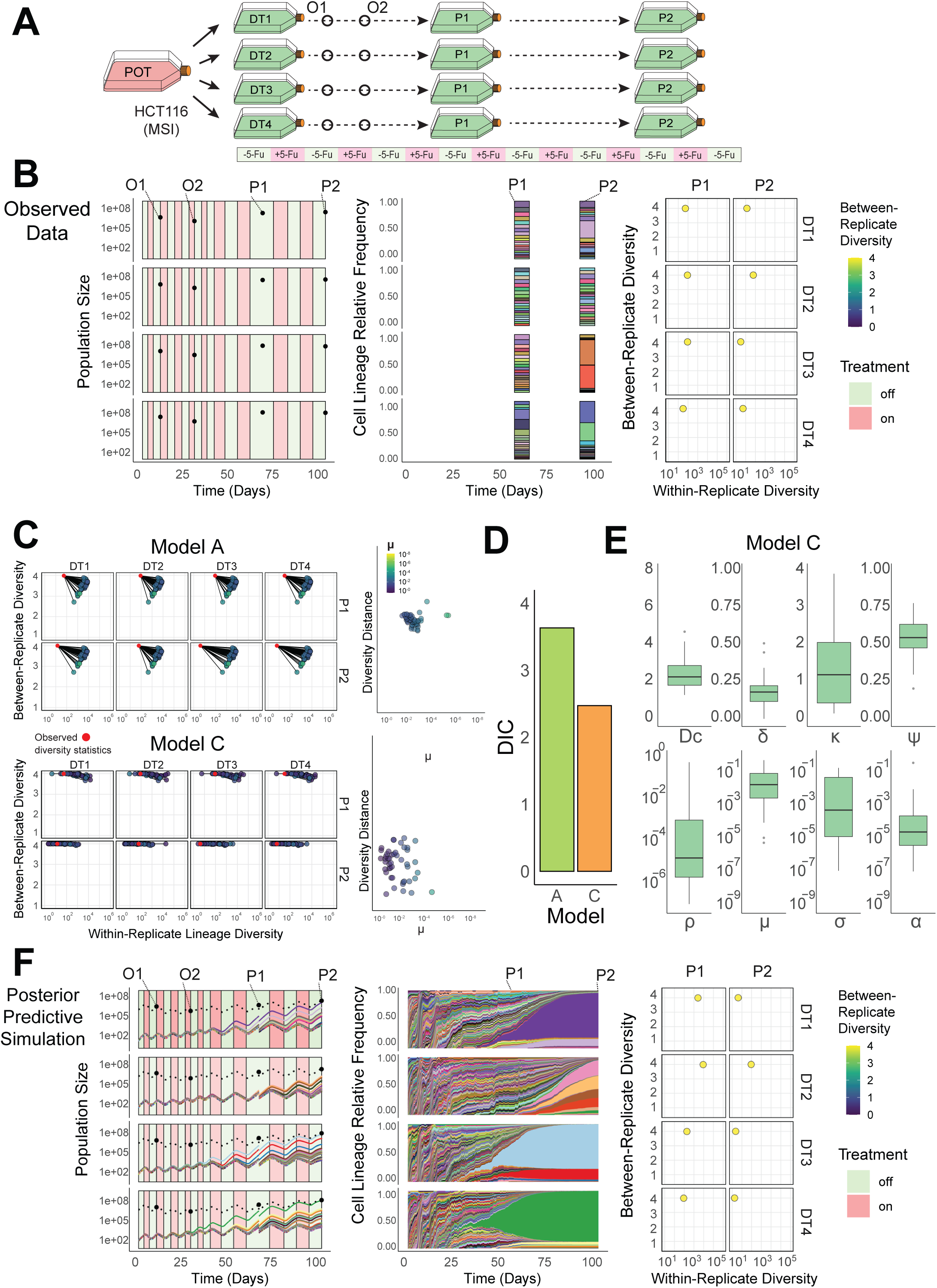
Quantification of the evolutionary dynamics during treatment in a barcoded colorectal cancer cell line (HCTbc). **A)** A simplified schematic of the experimental design used in a long-term evolutionary resistance assay. Barcoded HCT116 cells (HCTbc) were barcoded and expanded (POT) before being sampled into four replicate drug-treatment sub-populations (DT1-4) that were exposed to periodic chemotherapy treatment (5-fluoruracil: 5- Fu) and passaged at two timepoints (P1 & P2). **B)** Population size counts at four timepoints per-replicate (DT1-4), including two intermediate population size estimates (O1 & O2) and two Passage timepoints (P1 & P2) (left panel); top 20 sequenced barcode lineage relative frequencies at the two passage timepoints, where area colour denotes lineage identity (middle panel); lineage diversity statistics of the sequenced barcode distributions at the two passage timepoints (P1 and P2) for each replicate composed of within replicate lineage diversity and between-replicate diversity dissimilarity. **C)** Posterior predictive simulations for Model A (unidirectional transitions – top) and Model C (escape transitions – bottom) of the within and between-replicate diversity statistics (LHS) and the normalised diversity distance from the observed statistics (red points) to the simulated values (highlighted by the transition parameter μ). **D)** The Deviance Information Criterion (DIC), a measure of model fit, for Model A & Model C (lower values indicate higher model support) calculated using the posterior predictive distributions. **E)** The posterior distributions (n=50, 4 experimental replicates) for parameters in Model C (the model with the highest support). Boxplots show the median, the first and third quartiles, and whiskers 1.5x the interquartile range. Parameters: *ρ* - pre-existing fraction of resistance, *μ* – sensitive to resistant transition probability per cell division, *σ* – resistant to sensitive transition probability per cell division, *α* – resistant to escape transition probability per cell division, *ψ* – strength of the resistant phenotype, Dc – maximum strength of the drug, *κ* – accumulation/decay rate of the drug. **F)** A posterior predictive simulation using Model C.

In the barcoded cell line SW6bc, fitting our model A (‘unidirectional switching’) predicted that resistance was rare when the experiment began (*ρ* = 1.1×10^-4^ ± 2.3×10^-4^, mean ± s.d.) and that the phenotypic transition rate from sensitive to resistant was low (*μ* = 7.6×10^−7^ ± 1.5×10^-6^, mean ± s.d.). Resistant cells also experienced a significant probability of drug-induced death during treatment (*ψ* = 0.63 ± 0.01, mean ± s.d.) and the fitness penalty of resistant cells in the absence of treatment was moderately low (*δ* = 0.11 ± 0.08, mean ± s.d.) (Fig. 4E). These dynamics led to resistant lineages that were not dominant in the population until the end of the 2nd passage, as illustrated by the drop in within-replicate diversity between P1 and P2 (Fig. 4C - Top). The ‘unidirectional’ model was preferred over the more complicated ‘escape transition’ model, as supported by a lower DIC score (Fig.4D). The low frequency of resistance and its stability through cell divisions predicted by the model explain the high similarity of the lineage distributions (low between-replicate diversity values, Fig. 4B&F, right hand column): the correlation between lineage identity at the time of barcoding (t_0_) and the resistant phenotype remains largely unbroken through multiple rounds of treatment and population bottlenecks. The dynamics predicted by the modelling framework were consistent with a stable, pre-existing resistance mechanism that emerges gradually from a predominantly sensitive population in SW6bc.

In our second barcoded cell line, HCTbc, a few different unique lineages grew to dominate each replicate flask following treatment for two passages (Fig. 5B). To recover the population size changes and unique barcodes ‘winning’ across each replicate sub-population, we had to invoke our more complex model C, ‘escape transitions’ (Fig.5C & Fig.5D).

The model supported resistant cells incurring a fitness penalty (*δ* = 0.18 ± 0.08, mean ± s.d.) and there was a high probability of a transition from the sensitive to resistant phenotype per cell division (*μ* = 7.6×10^-2^ ± 0.11, mean ± s.d.). The probability which cells escaped the fitness penalty was lower (*α* = 2.5×10^−4^ ± 6.8×10^-4^, mean ± s.d.) (Fig. 5E). These dynamics led to sensitive cells having a moderate but equipotent capacity to switch to a slower growing resistant phenotype to survive initial rounds of treatment, followed by rarer ‘jackpot’ events where the lineage grew to dominate the population after escaping the fitness penalty (*α*). The lower probability of these jackpot events occurring within slower growing resistant cells also contributed to stochastic variation in diversity statistics between replicates (Fig. 5C – Bottom Panel), as observed in our sequenced barcode distributions.

This ‘two-stage’ evolutionary process predicted to be responsible for resistance in our barcoded HCTbc cell line is reminiscent of ‘persister’ dynamics observed elsewhere, whereby a population of cells with slower division rates initially survive treatment and provide a reservoir of cells within which additional adaptive changes can accrue (Álvarez-Arenas et al., 2019; Ramirez et al., 2016; Rehman et al., 2021; Windels et al., 2019).

In HCTbc, the model predicted the rate at which cells experience elevated death rates due to the change in effective drug concentration (*κ*) to be quicker compared to SW6bc. This difference meant the relative fitness benefit experienced by the resistant (vs sensitive) phenotype immediately following treatment was greater in HCTbc, consistent with the greater relative increases in the IC_50_ values for the HCTbc drug-treatment replicates (Supp. Fig. 7).

In summary, data from the SW6bc cell line were best explained by model A (unidirectional transitions), where resistance was rare and the transition rate from sensitive to resistant cells was low, consistent with a stable, pre-existing molecular driver of resistance. In contrast, data from the HCTbc cell line were better predicted by model C (escape transitions), where faster-growing, treatment-resistant cells emerged from a more common population of slower-growing, treatment-refractory cells.

### Functional characterisation of resistant cells verifies modelling inferences

We performed detailed molecular and functional characterisation of cells from our long-term resistance evolution experiment conditions. Whilst our framework does not infer the molecular mechanisms of resistance directly, it does make predictions for how the phenotypes (here, resistance to our chemotherapy 5-fluorouracil) should be distributed amongst a cell population.

To test the predictions generated from our modelling framework at the transcriptional level, we performed single-cell RNA sequencing (scRNA-seq) of the expanded parental population (POT), two fourth passage control (CO), two fourth passage drug-treatment (DT) replicates and two ‘drug-stop’ (DS) replicates (Fig. 6A). The DS replicates had only been subject to a single cycle of treatment and recovery before being harvested and hence primarily exhibited the phenotype of parental cells when exposed to treatment. We could therefore compare DS and DT conditions to the parental (POT) samples simultaneously to isolate gene expression changes that were either: an ‘acute treatment response’ following exposure to the drug (changes shared between DS and DT comparisons); a ‘sensitive treatment response’ in genes that were only found differentially expressed when the ancestral population was exposed to treatment (differentially expressed in DS but not DT comparisons); and a ‘resistance treatment response’ for genes differentially expressed in the evolved resistant lines but not the sensitive parental cells (differentially expressed in DT but not DS comparisons). Given these comparisons, gene expression changes associated with resistance can fall into two categories: either those that are found solely differentially expressed in the drug-treatment replicates (‘resistance treatment response’) or those whose differing expression in response to acute treatment exposure in the primarily sensitive ancestral population has been lost during the evolution of resistance in the drug-treatment replicates (‘sensitive treatment response’).

**Fig. 6.**
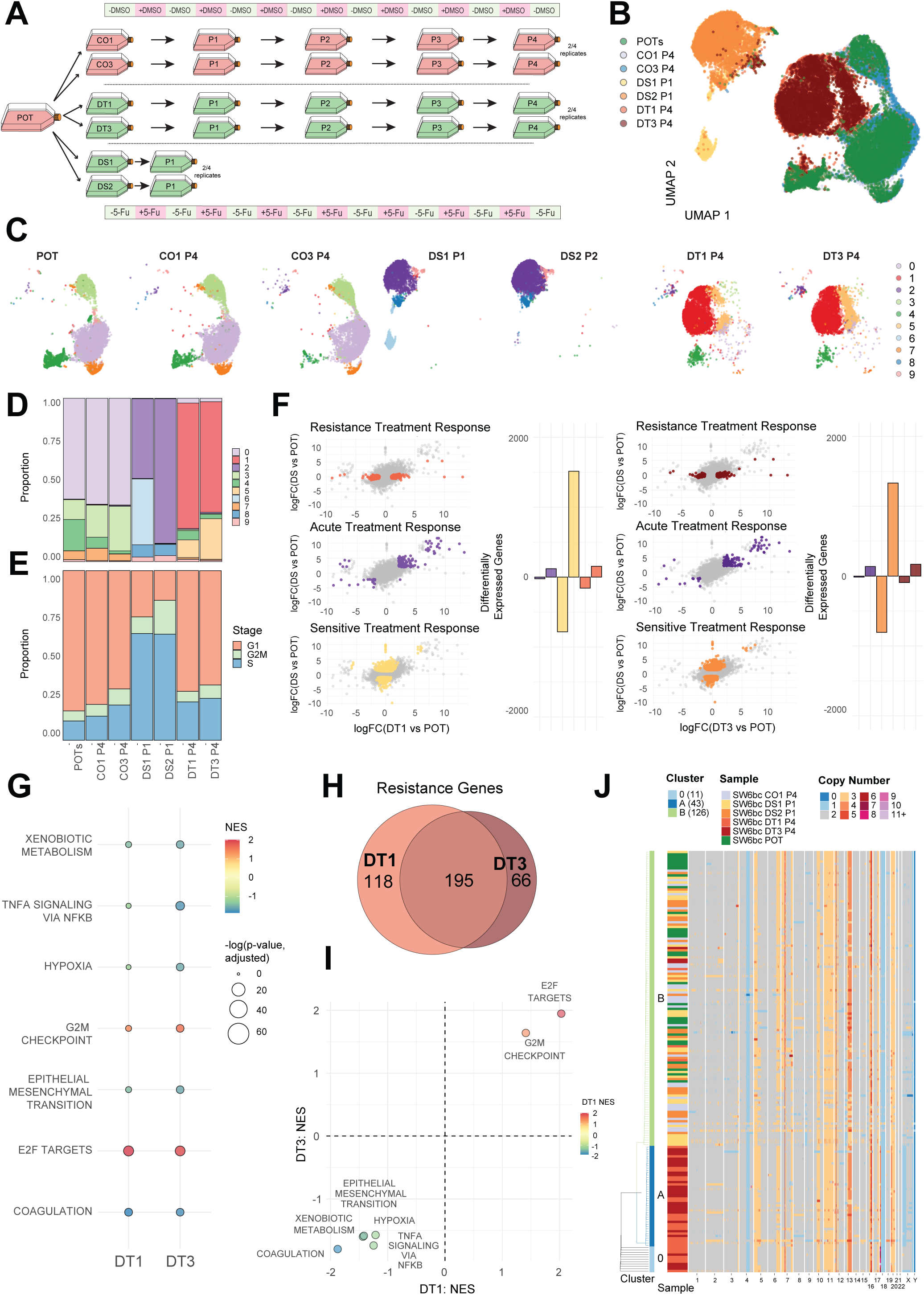
Functional characterisation of resistance in SW6bc. **A)** Schematic of sample design and naming schemes from the long-term resistance evolution experiment using barcoded SW620 colorectal cancer cells. Periods of on/off treatment (5-Fu and DMSO) are for illustrative purposes and are not to scale. **B)** UMAP dimensionality reduction of the scRNA-seq data from chosen sample replicates. **C)** Cluster assignment of UMAP results on a sample-by-sample basis highlighted by cluster identity. **D)** Cluster assignment proportions by cell sample. **E)** Cell cycle stage proportions by cell sample. **F)** Genes associated with replicate-specific differential expression (DE) identified via specific comparisons (coloured points per panel): Resistance Treatment Response - DE in DT but not DS; Acute Treatment Response - DE in DT and DS; Sensitive Treatment Response - DE in DS but not DT. Comparisons are performed for each drug treatment replicate (DT1 & DT3) and the number of significantly DE genes in each group are also shown per comparison. Significantly differentially expressed genes were identified using a quasi-likelihood F-test (QLF test) in edgeR. **G)** Gene set enrichment analysis (GSEA) for each drug-treatment replicate (DT1 and DT3) for genes identified as ‘Resistance’ genes in each replicate. Only gene sets that were found to be significant (adjusted p-value < 0.05) in one of the two replicates are shown. Point colour denotes normalised enrichment score (NES). Adjusted p-values were computed using the multilevel adaptive algorithm implemented in fgseaMultilevel. **H)** The number of shared/unique Resistance genes between each drug-treatment replicate. **I)** ‘Resistance’ genes GSEA NES comparison between DT1 and DT3. **J)** A subset of single-cell copy number profiles from SW6bc experimental replicates. Each row is a single cell and chromosome bin colour denotes copy number. Rows are highlighted by sample identity and cluster assignment.

In our barcoded SW6bc CRC cell line, our modelling framework predicted the existence of a pre-existing, stable molecular driver of resistance. The parental and control replicates exhibited highly similar expression profiles, supporting the experimental passaging and vehicle control treatment having a negligible effect on gene expression (Fig. 6B&C). The drug-treatment replicates also exhibited remarkably similar expression profiles with each other, despite several months of chemotherapy in isolated experimental replicates (Fig. 6C). Of the differentially expressed genes associated with resistance (Fig. 6F - ‘resistance treatment response’ signature), the majority were shared between the two sequenced replicates (Fig. 6H). Furthermore, the shared nature of gene expression was also present when comparing gene signature hallmarks, with high concordance in the normalised enrichment scores (NES) (Fig. 6G&I). Most gene expression differences across all comparisons in SW6bc were those unique to the DS replicates (Fig. 6F - ‘sensitive treatment response’ signature). Given DT and DS replicates have both been recently exposed to the chemotherapy 5-Fu, these findings suggest that evolution of resistance in SW6bc leads to cells that more closely resemble their untreated ancestors despite ongoing treatment. Overall, the scRNA-seq results described here are consistent with selection for a shared resistant phenotype across replicates, as predicted by our modelling results.

Given the putative chromosomal instability in this cell-line, we also looked for evidence supporting the selection of a rare pre-existing sub-population with a unique pattern of copy number alterations (CNAs) by measuring CNAs in individual cells. We performed single-cell shallow whole-genome sequencing via ‘Direct Library Preparation’ (DLP) on different cells from the same experimental conditions that were characterised via scRNA-seq (Zahn et al., 2017). The copy-number profiles supported ongoing chromosomal instability in SW6bc, with small sub-clonal changes found across all samples (Fig. 6J). Consensus copy number profiles derived using all cells within a sample identified additional changes shared by only the drug-treatment replicates (Supp. Fig. 8), consistent with the repeated selection of a shared ancestral lineage (as seen in the lineage tracing and transcriptomic data). Clustering based on the single-cell copy number profiles grouped most cells from each drug-treatment replicate (DT1 & DT3) together, also consistent with the expansion of a shared ancestral sub-clone (Fig. 6J).

In our second barcoded CRC cell line, HCTbc, we performed scRNA-seq on replicates with the same sample design as our previous cell-line (Fig. 7A). Whilst the high concordance between POT and CO replicates again supported a minimal impact of sampling and vehicle control treatment, the two replicates’ drug-treatment cells exhibited markedly distinct expression profiles (as represented via dimensionality reduction; UMAP - Fig. 7B&C). Also, in contrast to SW6bc, the DS replicates were enriched for the G1 cell cycle stage, consistent with elevated cell-cycle arrest in parental cells following a single round of treatment.

**Fig. 7.**
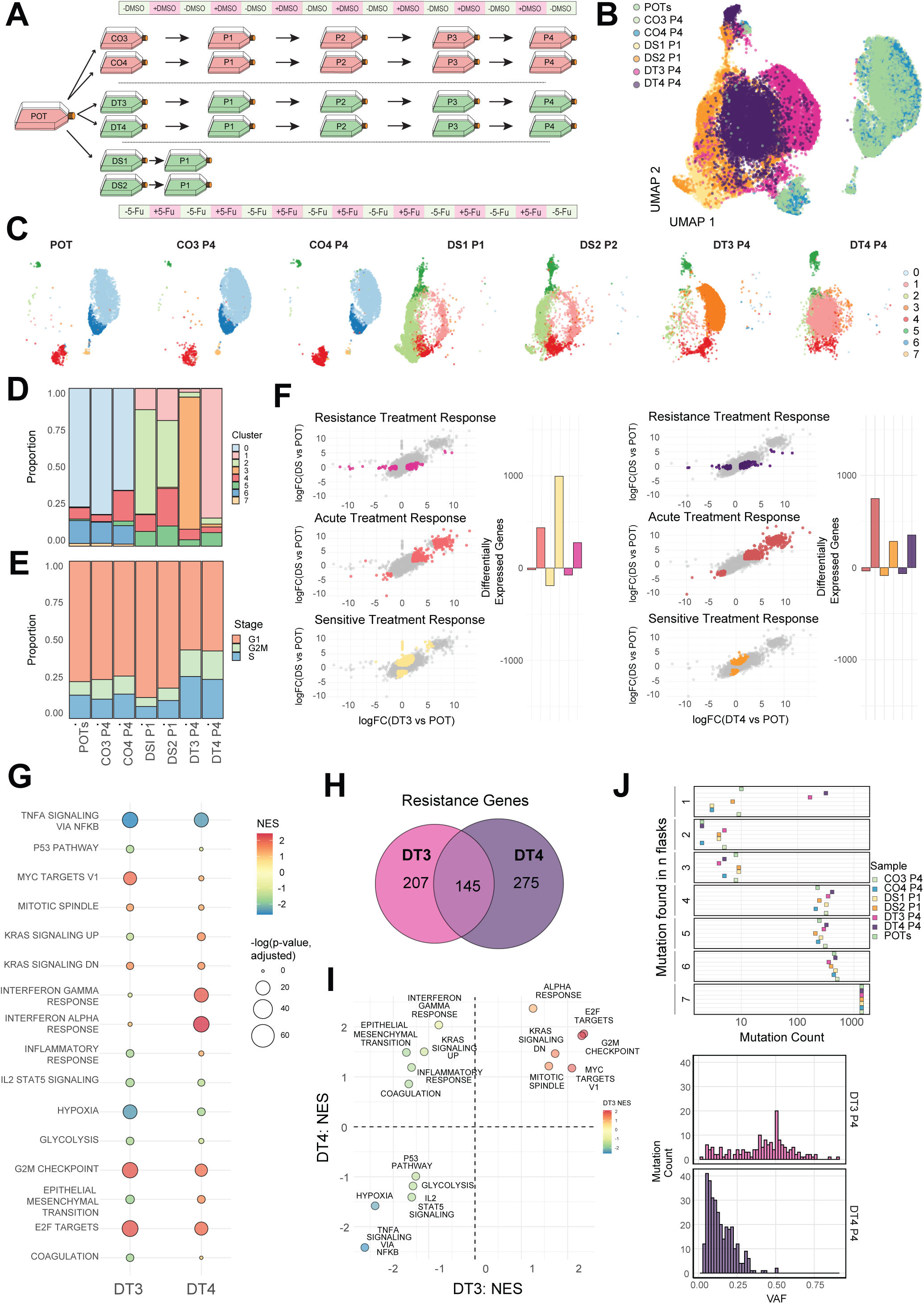
Functional characterisation of resistance in HCTbc. **A)** Schematic of sample design and naming schemes from the long-term resistance evolution experiment using barcoded HCT116 colorectal cancer cells. Periods of on/off treatment (5-Fu and DMSO) are for illustrative purposes and are not to scale. **B)** UMAP dimensionality reduction of the scRNA-seq data from chosen sample replicates. **C)** Cluster assignment of UMAP results on a sample-by-sample basis highlighted by cluster identity. **D)** Cluster assignment proportions by cell sample. **E)** Cell cycle stage proportions by cell sample. **F)** Genes associated with replicate-specific differential expression (DE) identified via specific comparisons (coloured points per panel): Resistance Treatment Response - DE in DT but not DS; Acute Treatment Response - DE in DT and DS; Sensitive Treatment Response - DE in DS but not DT. Comparisons are performed for each drug treatment replicate (DT3 & DT4) and the number of significantly DE genes in each group are also shown per comparison. Significantly differentially expressed genes were identified using a quasi-likelihood F-test (QLF test) in edgeR. **G)** Gene set enrichment analysis (GSEA) for each drug-treatment replicate (DT3 and DT4) for genes identified as ‘Resistance’ genes in each replicate. Only gene sets that were found to be significant (adjusted p-value < 0.05) in one of the two replicates are shown. Point colour denotes normalised enrichment score (NES). Adjusted p-values were computed using the multilevel adaptive algorithm implemented in fgseaMultilevel. **H)** The number of shared/unique resistance genes between each drug-treatment replicate. **I)** ‘Resistance’ genes GSEA NES comparison between DT3 and DT4 **J)** Mutations found in scRNA-seq data grouped by the number of replicate flasks any mutation was found in and VAF distributions of mutations found in scRNA-seq data unique to the two drug-treatment replicates.

In one of the two drug-treatment replicates characterised (DT3), the majority of differentially expressed genes across our joint comparisons were those that were only found in the parental cells’ response to treatment (DS - ‘acute treatment response’, Fig. 7F). However, unlike SW6bc, the other replicate (DT4) shared a higher proportion of differentially expressed genes with the parental population’s response to treatment (POT vs DS *and* POT vs DT - ‘acute treatment response’, Fig. 7F). These suggest that one of the two replicates (DT3) now more closely resembles the untreated ancestral phenotype (POT) despite ongoing treatment. Alongside most resistance associated genes being unique to each drug-treatment replicate (Fig. 7H), these results supported the emergence of unique resistant phenotypes in HCTbc. Interestingly, in contrast to SW6bc, this non-repeatability was observable when comparing hallmark gene sets: whilst some pathways showed concordance in the magnitude and direction of enrichment, some pathways showed different directions between the two replicates, including EMT genes and genes associated with elevated KRAS signalling (both showed elevated enrichment in DT4 but depletion in DT3 - Fig. 7I).

As HCTbc was characterised by microsatellite instability, we investigated the clonal dynamics associated with resistance evolution by comparing the mutational landscape of the cells (called from scRNA-seq data). We compared the number of mutations found in *n* of the total sequenced replicate flasks (Fig. 7J). Many mutations were found that were unique to each drug-treatment replicate (Fig. 7J- Found in 1). Comparison of the VAF distributions supported more complete dominance of the ‘jackpot’ lineage (predicted by the modelling) in the DT3 replicate, where a large number of mutations had reached clonality (VAF ∼0.5), whereas in DT4 the stochastic nature of the successful lineage emergence had either led to clonal interference mitigating clonal dominance, or incomplete dominance of the faster proliferating resistant lineage in the population. This latter hypothesis was supported by a ‘fitness assay’: single cell isolated clones derived from DT4 were significantly slower growing than those from DT3 (Supp. Fig 9), and the transcriptional results which also supported DT4 more closely resembling the phenotype of the parental cells recently exposed to treatment (the DS condition).

Our modelling framework predicted evolution during treatment in HCTbc occurred via a two-step process: treatment refractory cells that exhibited a fitness penalty (realised as slower net growth rates) initially survived treatment (the ‘resistant’ phenotype), and cells that escape this fitness penalty but remained resistant emerged stochastically from this population as treatment continued (the ‘escape’ phenotype). We reasoned that our original scRNA-seq sampling design alone was limited in its ability to distinguish these dynamics from other evolutionary scenarios. Specifically, because the transition to ‘escaped’ was predicted to have occurred between seeding and the first passage, waiting for the cells to fill the flask means observations were limited to cells that had emerged following the second event.

Motivated by this, we performed an additional drug-treatment experiment where we exposed parental barcoded cells (HCTbc POT) to periodic chemotherapy using the treatment regime adopted during our original experiment. We waited until colonies of cells appeared that could readily proliferate despite ongoing treatment, and we observed that when this occurred the dividing cells were smaller than the majority of those that had initially survived treatment (Supp.Fig.10A). We therefore predicted we could use size as a proxy for cells occurring pre- and post- the ‘escape’ event and isolated single small or large cells before performing scRNA- seq. We also sequenced (scRNA-seq) single cells from the original HCTbc experimental conditions (Fig. 6A) to validate these new comparisons.

To identify differentially expressed genes relative to the parental (POT) cells but either shared with or independent of the parental population’s response to treatment (DS), we performed analogous comparisons to those discussed previously to identify a signature of: ‘large cell treatment response’ (changes associated with large cells but not DS replicates); ‘small cell treatment response’ (changes associated with small cells but not DS replicates); ‘acute treatment response’ (changes associated with either large or small cells and DS replicates); and ‘sensitive treatment response’ (changes associated with the DS replicates but not either large or small cells). Whilst the large cells exhibited the highest number of differentially expressed genes in response to treatment vs the DS replicates, performing these comparisons with the small cells revealed a high number of genes in the ‘sensitive treatment response’ group (Supp. Fig. 10B). Finding the majority of differentially expressed genes unique to the DS replicates was consistent with the emergence of a readily proliferating small cell phenotype that more closely resembles the ancestral (POT) cells. Correlations between average cell expression profiles also support a small cell phenotype that more closely resembles the parental cells, whereas the large cells appear transcriptionally distinct (Supp. Fig. 10C). Interestingly, gene set enrichment analysis on genes that were differentially expressed in the large cells (relative to the POT replicates) showed down regulation of pathways that were elevated in the HCTbc drug-treatment replicates (Supp. Fig. 10D). In contrast, these pathways were up-regulated in the small cell phenotype (Supp. Fig. 10E), consistent with the drug-treatment replicates. These findings supported two distinct phenotypes existing following treatment in HCTbc, as predicted by our model. Whilst we cannot be certain that the presence of large and small cell phenotypes are indicative of a process whereby the latter emerges from the former, the existence of distinct colonies who have gene expression changes that more closely resemble resistant cells suggest rare, stochastic changes render cells able to proliferate despite the presence of treatment.

In summary, our functional characterisation of cells derived from the same experiment used to infer evolutionary dynamics from population size data and lineage distributions supported the predictions made by our modelling framework. In SW6bc (a CIN barcoded CRC cell line) a simple model of a stable, pre-existing molecular driver of resistance was most likely. In contrast, in HCTbc (a MSI barcoded CRC cell line), a more complicated two-step model was required to explain the data.

## Discussion

Here, we report a framework for combining population size changes through time with lineage distributions to provide quantitative insight into the dynamics of resistance evolution. Whilst some dynamics can be inferred from a population’s changing size in response to treatment, we show that explicitly incorporating cell lineage relationship information can markedly increase the ability to identify phenotypic behaviours; for example, rates of phenotypic switching, the strength of treatment selection and phenotype-dependent fitness penalties can be measured simultaneously without having to perform separate, technically arduous assays for each behaviour of interest. These findings have broader ramifications for approaches that aim to predict phenotypic compositions and behaviours of cancer cell populations solely from population size measurements. They suggest that metrics such as tumour burden alone may be insufficient to predict the change in sensitive and resistant populations over time, for example, and that measurements of cell lineage relationship changes through treatment are a necessary addition.

Although the power of our modelling framework is limited to scenarios where populations initially respond to treatment or where a portion of measurable lineage diversity is retained for the observable duration, we argue there are few dynamics to learn when resistance is complete from the start. Additionally, newer barcoding technologies allow for the ongoing labelling of lineage identities, precluding the complete loss of measurable lineage diversity through treatment (Bhattarai-kline et al., 2022; Yang et al., 2022). Here, we used a simple re-barcoding step to explore where these barcodes might provide the greatest advantage. We found that clonal sweeps that occur following the initial crash in diversity enabled the re-barcoding step to recover the transition rate from slow growing and tolerant to resistant and proliferating. We believe the combination of evolving barcodes that are also accessible in transcriptional readouts with the modelling framework described here will further facilitate the easy experimental identification of routes to resistance.

There are important differences between clinical and experimental data that could limit the direct application of our framework to patient samples. Some of the statistical power to distinguish between possible evolutionary parameter combinations comes from the ‘evolution in parallel’ information provided by experiments where closely related populations (replicates) are subjected to the same selection pressure synchronously. In patients, no equivalent replication exists. Additionally, our models assume that spatial factors do not impact evolutionary dynamics, whereas, in reality, the 3D structure of a tumour could impose constraints that cause deviations from our assumption of a well-mixed population. There are also practical challenges in generating the time-series tissue samples required to measure the cell lineage compositions of a patient’s tumour. Nonetheless, translation to patient samples should be possible because mutations in somatic cells can provide naturally occurring barcodes that permit retrospective lineage tracing (Fabre et al., 2022; Gabbutt et al., 2022a; Warner et al., 2024). Given joint observations of a patient’s total tumour size, a readout of lineage identities and known treatment timings, our approach could be implemented to infer the dynamics of resistance evolution in a clinical setting with greater resolution than using burden or lineage observations in isolation.

Our modelling framework focuses on the transition rates between phenotypes opposed to the class of molecular change responsible for the difference, i.e. if they are genetic or non-genetic. As such, we do not restrict ourselves towards finding one class over the other, but instead can exclude genetic mechanisms only if the transition rates are high enough (as was the case for the transition from the sensitive to slower growing, resistant phenotype in our barcoded HCTbc cell line). However, we do note that for very slow transitions, distinguishing a stable epigenetic mechanism from a genetic one is not possible. Indeed, positive selection for an epigenetic modifier would resemble that of a genetic mutation if it persisted within a cell lineage for many divisions. We were initially surprised by the framework’s inability to recover the reversion switching rate from a resistant to sensitive phenotype. It appears that any switches to a phenotype that experience negative selection when the environment changes are difficult to detect. We note that it is possible that only under controlled experimental single-cell isolation this reversion behaviour can be accurately quantified. Even in cases where populations are exposed to the same pressure of treatment after a recovery period (often coined a ‘treatment holiday’), movement in the population’s mean phenotype to sensitivity could also be due to differences in phenotype specific growth rates. Frequency-dependence in growth and switching probabilities may also impact the ability to recover these evolutionary parameters of interest.

Our framework can identify numerous modes of resistance evolution to a given treatment. For example, it can be used experimentally to easily identify whether resistance is the product of a stable, heritable change or more transient non-genetic switching phenomena. Such an approach could help guide the development of treatments by enabling a focus on putative resistance mechanisms early in the process. We believe the explicit incorporation of lineage information will increase the rate and accuracy of resistance mechanism identification, permitting better access to novel cancer vulnerabilities.

## Methods

### *In vitro* long-term experimental resistance evolution

#### Cell culture

Two colorectal cancer cell-lines HCT116 (ATCC CCL-247™) and SW620 (ATCC CCL-227™) were used for all data generating experiments in this project. Cells were grown in ‘standard conditions’ which were as follows: growth medium, consisting of high glucose DMEM supplemented with 10% Foetal Bovine Serum (FBS) (Gibco, A5256701) and 2% Penicillin-Streptomycin (Gibco, 15140122) (from hereon in: ‘full growth medium’, unless stated otherwise). Cells were grown in T-175 vented flasks (Corning, 431080) for the long-term drug-treatment experiment with 35mL of full growth medium. Flasks were grown at 37°C, 5% CO^2^ and 95% relative humidity. Cells were cryopreserved using a slow freezing technique. Briefly, cells were resuspended in freezing medium consisting of 90% FBS and 10% dimethyl sulfoxide (DMSO) (Sigma-Aldrich, D2650). The cell suspension was aliquoted into cryovials, which were then placed in Mr. Frosty^TM^ freezing containers (Thermo Fisher Scientific, 5100-0001) before being stored at −80°C overnight. After 24 hours, the vials were transferred to liquid nitrogen for long-term storage. Cell line identities were confirmed using STR analysis at the beginning of the experiment and were tested regularly for mycoplasma infection.

#### Lineage tracing

The ClonTracer library was a gift from Frank Stegmeier (Addgene #67267). The library was expanded in-house and optimisation steps also undertaken according to the original ClonTracer protocol such that the multiplicity of infection (m.o.i) of cells was estimated to be 0.1, minimising the number of multiple integrations (Bhang et al., 2015). In addition, optimisation experiments were performed for each cell line (HCT116 and SW620) such that we could confidently barcode approximately 1×10^6^ cells (per cell line) following infection and puromycin selection. The infection and selection steps for the long-term resistance experiment were split over 3×150mm (Corning, 430599) plates to ensure there was sufficient room for population expansion following barcoding without the need for additional passaging events. Given our framework uses the lineage distributions enabled by these barcodes to estimate the evolutionary dynamics, we performed single cell colony expansion and sequencing of some representative clones to make sure that most cells did contain a single barcode: cells were diluted to a concentration of either 100 cells/200μl (One control well of a TC-treated 96-well adherent plate (Corning, CLS3596)) or 1 cell/200μl (remaining wells of a 96-well plate) of conditioned growth media (50% standard media, 50% media from cells that were 80% confluent, filtered in 0.45μm size filter). This process was repeated for a total of 5x 96-well plates per cell line. After 7 days, wells were checked for visible single colonies. After another 21 days, colonies were harvested and the barcodes in each expanded colony sequenced (details provided in DNA Extraction, Barcode Amplification and Barcode Sequencing).

#### Long-term resistance evolution assay

Following barcoding of colorectal cancer cell lines with the ClonTracer genetic lineage tracing system, we expanded the initial population to ensure that each lineage was represented multiple times. We froze vials of this expanded population, POT, and seeded 12 replicate sub-populations of 1×10^6^ cells each for 4 replicates across 3 conditions: vehicle control, CO, drug-stop, DS, and drug-treatment, DT. This design was repeated in parallel for each barcoded cell line (HCTbc and SW6bc). Control replicates were treated with periodic vehicle control (DMSO) for 4 passages, drug-treatment replicates with IC_50_ values of the chemotherapeutic 5-fluoruracil (5-Fu) (Selleckchem, S1209) for 4 passages and drug-stop with a single period of IC_50_ 5-Fu treatment then a single recovery period before harvesting and freezing (details provided in Drug Inhibition Curves methods section). All flasks were passaged before 80% confluency was reached. When passaging, 1×10^6^ cells were seeded into the subsequent replicate flask and the sample’s remaining cells were split into two vials and frozen. Frozen cells were stored for future DNA extraction, barcode amplification and additional functional assays.

#### DNA extraction, barcode amplification and barcode sequencing

To extract genomic DNA for barcode amplification, cells were defrosted and a volume containing approximately 1×10^6^ cells was spun down into a pellet and re-suspended in 200µL of Phosphate Buffered Saline (PBS) (Gibco, 10010023). DNA was then extracted according to the DNeasy Blood & Tissue Kit (QIAGEN, 69506) protocol. DNA was eluted in DNase and RNase-Free PCR grade water. To quantify the concentration of DNA, the eluted volume was vortexed and then 1µL taken and measured using a Qubit 4 Fluorometer (Invitrogen, Q33226) according to the manufacturer’s protocol. DNA concentrations were recorded and the DNA frozen at −20°C until future processing. Following PCR steps, a magnetic bead based clean-up system was used for the purification of amplified barcode DNA (CleanNGS – CleanNA, CNGS-0050).

To amplify the barcodes from cell DNA, two universal amplification primers targeted the universal sequences flanking the semi-random barcode sequence in the ClonTracer construct directly (CT_UV_FWD: ACTGACTGCAGTCTGAGTCTGACAG and CT_UV_REV: CTAGCACTAGCATAGAGTGCGTAGCT). A bespoke adapter ligation protocol was optimised that used unique dual-indexes (IDT Illumina™ TruSeq UD Indexes 96) (Illumina, 20040870) - each sample was identified via its own unique forward and reverse indexes. This approach minimised index hopping, a common problem for low complexity libraries sequenced on newer Illumina patterned flow cell systems (Costello et al., 2018). Final libraries were quantified using Agilent HS D1000 screentapes (Agilent, 5067-5584) on an Agilent Tapestation 4200 system and sequenced on the Illumina Novaseq 6000 system using Novaseq S2 PE50 flow cells. Due to the low-diversity amplicon library, 15% of Phix control library was used.

### Phenotypic and molecular characterisation of phenotypes

#### Drug inhibition curves

The IC_50_ values of each cell line were measured as follows: Cells were trypsinised into a single-cell solution and cell viability was confirmed to be > 80% using the Countess™ II Automated Cell Counter (Thermo Fisher Scientific). Cells were then seeded into a TC-treated 96-well adherent plate (Corning, CLS3596) (8000 cells/well for HCT116 and 10000 cells/well for SW620) in 200µL of full growth medium. After 1 day (+24hrs from start), full growth medium + given drug concentrations were added to the respective wells. After three additional days (+96hrs from start), CellTiter-Glo ® Reagent (Promega Ltd, G9683) was thawed and 50µL of reagent was added to all wells. Plates were placed in the incubator at standard conditions for 15 minutes before luminescent readings were taken using a PHERAstar FSX plate reader (BMG Labtech, Germany). The results were passed to a custom R script to calculate the dose-response curve (Ritz et al., 2015).

#### Single cell sorting for growth rate assays, scWGS and size-sorted scRNA

Barcoded HCT116 (HCTbc) and SW620 (SW6bc) cells from the long-term resistance evolution assay were single cell sorted for colony growth assays (HCTbc) and single-cell whole genome sequencing (scWGS, SW6bc). Vials were thawed and expanded briefly (<3 days) from the respective replicates. A cell count of >1×10^6^ and viability of >80% ensured there was no stringent bottleneck prior to characterisation. Single cell solutions were obtained by resuspending trypsinised cells in PBS and single cells were sorted into single wells using the cellenONE ® single cell sorting platform (Cellenion). For the colony growth assays, cells were sorted into wells of a 96-well plate which contained conditioned growth media (50% standard media, 50% media from cells that were 80% confluent, filtered in 0.45μm size filter). For scWGS, cells were sorted into wells that contained a pre-distributed lysis buffer (Laks et al., 2019). For scRNA-seq using the cellenCHIP 384-3’RNA Seq Kit, single cells were sorted directly into the chip: large cells were chosen as those with a diameter >36µm and an elongation factor <1.50 whilst small cells were chosen with a diameter filter of <20µm and an elongation factor <1.50.

#### Growth rate assays (HCTbc)

To measure the growth rate of single cell colonies derived from 2 control (CO) and 2 drug-treatment (DT) HCTbc replicates, single cell sorted plates were kept in an Incucyte ® S3 Live-Cell Analysis System, and confluence readings were recorded every 12 hours for 38 days. As most cells were sorted near the edge of the well, any wells that contained a cell sorted at the centre were excluded to control for well-position dependent differences in growth kinetics. After the full expansion duration, confluence readings were exported, and growth rates extracted via custom R scripts by fitting a linear regression model to the log-confluence values.

#### Single-cell whole-genome sequencing (scWGS - SW6bc)

For whole genome sequencing of single cells from our barcoded SW6bc cells, we adopted the published protocol in Laks et al. (2019). First, 1ul each of 384 unique i5 indexing primer at a concentration of 4uM were dispensed into empty 384-well plates and air-dried. 1ul lysis buffer, consisting of 86.2% DirectPCR Lysis Reagent (Cell) (Viagen, 301-C), 8.6% protease (Qiagen, 501-PK), 5.2% glycerol (Sigma, G5516) was added to each well and a single SW620 cell was sorted into each well using the CellenOne F1.4 system (Cellenion). Following centrifugation of the plate at 3000G for 5 min and overnight incubation at +4°C cell lysis was performed by incubating the plate at 50°C for 1h and 70°C for 15min. Tagmentation of genomic DNA was carried out by adding 1.8ul tagmentation mix (Diagenode), consisting of 1.775ul tagmentation buffer (Diagenode, C01019043), 0.0165ul Tween (Sigma, P6585), 0.00875ul Tn5 loaded (Diagenode, C01070012), and incubating at 55°C for 10min. Subsequently, the reactions were neutralised by adding 1ul neutralisation mix (0.5ul Qiagen protease, 0.01ul Tween and 0.049ul ddH2O) and incubating the plate at 50°C for 15min and 70°C for 10min. Finally, PCR amplification was performed after adding 3.8ul master mix (3.78ul 2x NEB Ultra II Q5 master mix (NEB, M0544L), 1mM i7 indexing primer (500nM final, IDT, 10005974)) with the following cycling parameters: Gap-filling at 72°C for 5min, initial denaturation at 98°C for 30s followed by 10 cycles of 98°C for 30s and 65°C for 75s and final elongation at 65°C for 5 min. PCR products were pooled and purified using a Zymo DCC5 spin column (D4014) and then subjected to Exonuclease I digestion (NEB) and 1x Ampure bead purification. Final libraries were quantified using Agilent HS D1000 screentapes on an Agilent Tapestation 4200 system and sequenced on the Illumina Novaseq 6000 system using Novaseq S2 PE50 flow cells.

#### scRNA sequencing (HCTbc & SW6bc)

To perform scRNA-seq, cell samples were thawed (1×10^6^ cells/sample vial) and cell viability was confirmed to be > 80% using the Countess™ II Automated Cell (Thermo Fisher Scientific). Samples were then treated for one additional round of drug-treatment (IC_50_ 5-Fu for DS/DT conditions), vehicle control (CO condition) or grown in standard culture conditions (full growth medium – POT condition) for 3 days followed by 3 days of standard culture conditions for all conditions. Cells were trypsinised and cell viability confirmed to be > 80% and a fraction were re-suspended in PBS for an estimated number of 6000 captured cells per sample. The single cell suspension was loaded on a Chromium Single Cell 3′ Chip C, followed by Chip D (10X Genomics). A single-cell gel bead-in-emulsion was generated using the Chromium Single Cell DNA kit and the Chromium Controller. The scRNA-seq library was prepared, and the concentration and quantity of the complete library was confirmed using the TapeStation High Sensitivity Screen Tape assay D1000 (Agilent) and a High-Sensitive Qubit™ dsDNA Kit (Life Technologies). Samples were normalised, pooled, and sequenced on an Illumina NovaSeq 6000 according to standard 10X Genomics recommendations at a median depth of at least 50K read pairs per cell. All samples from the same cell line were prepared on the same day and all samples from each cell-line were sequenced on the same flow cell to minimise technical biases.

For the ‘size-sorted’ samples, we re-sequenced experimental conditions (POT, CO, DS, DT) from the original experiment (barcoded cell line HCTbc) alongside ‘large’ and ‘small’ cells following 7 weeks of treatment with IC_50_ values of 5-fluoruracil, and scRNA-seq was performed using the cellenCHIP 384-3’RNA-Seq Kit (Cellenion, CTR-5016). This kit includes the cellenCHIP 384, a nanowell array containing oligo-dT primers, unique cell barcodes (CB), and unique molecular identifiers (UMIs) for cDNA generation. The barcode oligos in the wells were rehydrated with Lysis and RT buffers followed by cell isolation and dispensing using the cellenONE (Cellenion). Reverse transcription was performed at 42°C for 90 minutes. The resulting cDNA was pooled by inverting the cellenCHIP 384 and centrifuging it into a recovery funnel for collection into microcentrifuge tubes. cDNA quantification was conducted using the High-Sensitive Qubit™ dsDNA Kit (Life Technologies), and amplification was carried out for up to 18 PCR cycles. The amplified cDNA was used to generate Illumina sequencing libraries, which were confirmed using the TapeStation High Sensitivity Screen Tape assay D1000 (Agilent).

### Computational simulations

To model resistance evolution, we employed two separate simulation approaches. In the first, we designed a hybrid model that simulates the total population sizes in distinct phenotypic compartments during periodic treatment. This model switched between a stochastic (jump process) and deterministic (ordinary differential equation) model when a population threshold was crossed to maintain stochastic dynamics experienced at low population sizes. Depending on the model in question, the simulation tracks 2 (unidirectional transitions (Model A) and bi-directional transitions (Model B)) or 3 (escape transitions (Model C)) phenotypic compartments that differ in their behaviour when exposed to treatment. This simulation is restricted to generating the total cell population size of each phenotypic compartment and does not record cell lineage relationships.

Our second simulation approach was a fully stochastic agent-based model simulated via a rejection-kinetic Monte Carlo algorithm. Here, individual cells were assigned lineage identities at the start of the simulation and the model output included total population sizes per phenotypic compartment (as in the hybrid model) but also lineage size distributions.

We adopted a Bayesian approach for parameter inference: specifically, the more computational expedient hybrid model was used for the initial generations of approximate Bayesian computation (ABC), where observed data were compared to simulated population trajectories. Later generations simulated lineage distributions with the fully stochastic model and the distance was computed between observed and simulated lineage diversity statistics to generate the final posterior distribution.

All computational simulations were written in Julia (version 1.7.2) (Bezanson et al., 2017).

### Bioinformatic analysis

#### Read merging, barcode extraction and clustering

Sequenced FASTQs from demultiplexed amplified barcode samples were merged using NGmerge (Gaspar, 2018) in ‘stitch mode’ with the following options: ‘-m 14, -p 0.2, -z’. The barcodes were then extracted and clustered using ‘Bartender’ (Zhao *et al*., 2018) with the following options: (extract) ‘-q-p GACAG[30]AGCAG -m 2 -d b’; (cluster) ‘-c 10 -d 2 -z 5’. Due to issues with index-hopping encountered when previously sequencing lineage tracing marks without the use of non-redundant indexes, we adopted unique dual indexes (UDIs) for barcode sequencing. Despite this, we still observed patterns suggestive of index-hopping in some of our samples (albeit to a much lesser degree). For example, barcodes that were found at high frequency in one cell line would often be found in low (but proportional) frequencies in the alternate cell line, despite independent barcoding steps and therefore putatively independent lineage relationships. These patterns also only appeared when samples were sequenced on the same flow-cell, confirming their technical nature. We therefore created a list of ‘problem barcodes’ when these patterns were observed and excluded them when calculating lineage diversity statistics. Of note, as we chose diversity statistics that leveraged higher frequency lineages, these exclusions of low frequency ‘swaps’ will have minimal impact when calculating the diversity statistics.

#### scRNA-seq expression and mutation analysis

For each barcoded cell line processed - HCTbc and SW6bc - the raw sequencing data were processed using the Cell Ranger software (10X Genomics) to perform sample de-multiplexing, barcode processing, and single-cell 3’ gene counting. The cellranger count pipeline was used for each sample, aligning reads to the GRCh38-2020-A reference genome and generating feature-barcode matrices for further analysis. The resulting filtered feature-barcode matrices were imported into R for downstream analysis using the Seurat package (Hao *et al*., 2021). Cells expressing fewer than 500 genes and with fewer than 1000 RNA molecules detected, and cells with more than 20% mitochondrial genes expressed were excluded from further analysis to remove low-quality cells. To standardize cell numbers across samples and prevent biases due to variations in the number of single cells sequenced, 6000 cells per sample were used for scRNA-seq analysis (the sample with the lowest cell count had >6000 cells sequenced). The ‘DoubletFinder’ package was used to find and exclude putative doublets (McGinnis, Murrow and Gartner, 2019). We normalised expression with the SCTransform function (Seurat v4.10) and used the normalised expression matrix for all downstream analysis. We identified the K-nearest neighbour graph based on the PCA-reduced dimensions using the FindNeighbors function (Seurat v4.10). The number of PCA dimensions used for this analysis was set to the first 30 components, which were determined based on the elbow plot method to capture the majority of the variance in the data. We performed clustering using the FindClusters function in Seurat using resolution=0.2. The clusters were identified based on the Louvain algorithm applied to the K-nearest neighbour graph.

To call mutations from the scRNA-seq data, we used the output Cell Range BAMs with SComatic (Muyas *et al*., 2023) with the following settings - Splitting Alignment Files: n_trim = 5; max_nM = 5; max_NH = 1; Collecting Base Count Information: min_dp = 10; min_cc = 10; min_bq = 30; Detection of Somatic Mutations: min_cov = 10; min_ac_cells = 3; min_ac_reads = 4; max_cell_types = 7; min_cell_type = 4.

#### scWGS analysis

Demultiplexed dual-index fastq files were obtained for single cells undergoing the DLP+protocol and the data were used as input for the workflow automation pipeline designed for the DLP+ method (https://github.com/shahcompbio/single_cell_pipeline) using default settings. First, raw reads were adapter-trimmed using TrimGalore, mapped with BWA aln to the hg19 reference genome and deduplicated using picard MarkDuplicates. Subsequently, copy number calling was performed using the HMMcopy tool with reads segmented into non-overlapping 500kb genomic regions (Daniel Lai, 2017). Cells were required to have a minimum quality score of 95, a minimum of 250,000 reads and an S-phase probability of less than 0.1. Only bins considered “ideal” in all cells were used. A set number of cells was sub-sampled from each sample (n=50) then dimensionality reduction, clustering and consensus copy number calls were performed using the R package ‘signals’ (Funnell *et al*., 2022).

#### Differential Expression Analysis

Differential expression analysis was performed in R using the edgeR package (Robinson, McCarthy and Smyth, 2010). scRNA-seq data (raw RNA counts) was combined into pseudobulk expression count matrices for each replicate sample. Each barcoded cell line (HCTbc and SW6bc) was analysed separately. Negative-binomial dispersion was estimated using the ‘estimateDisp’ function, and then fit the GLM to the count data with the ‘glmQLFit’ and ‘glpQLFTest’ to derive log-fold changes and FDR values for each gene. When identifying differentially expressed genes that were associated with a specific sample comparison, we selected those with a logFC > 2.0 and FDR < 0.05. When choosing genes that were not differentially expressed given the comparison between two samples, we chose genes with a logFC < 1.0 and FDR > 0.10. These subsets of genes were used to perform gene ontology (GO) or KEGG pathway analysis (Ritchie *et al*., 2015). For gene set enrichment analysis (GSEA) (Korotkevich *et al*., 2016), we created custom ranks based on the combinations of sample comparisons in question: if looking for genes associated with each comparison, we would take the product of both logFC values. If looking for genes whose expression was associated with one sample comparison but not the other, we generate a score with logFC_1 x (1 / (|logFC_2|+1)), where logFC_1 is the log fold change in expression in comparison ‘1’, the comparison where we are interested in differentially expressed genes. This custom score simultaneously penalises genes who have a high differential expression in comparison ‘2’ (logFC_2).

## References

Acar, A., Nichol, D., Fernandez-Mateos, J., Cresswell, G.D., Barozzi, I., Hong, S.P., Trahearn, N., Spiteri, I., Stubbs, M., Burke, R., Stewart, A., Caravagna, G., Werner, B., Vlachogiannis, G., Maley, C.C., Magnani, L., Valeri, N., Banerji, U., Sottoriva, A., 2020. Exploiting evolutionary steering to induce collateral drug sensitivity in cancer. Nature Communications 11, 1–14. 10.1038/s41467-020-15596-z

Acar, M., Mettetal, J.T., van Oudenaarden, A., 2008. Stochastic switching as a survival strategy in fluctuating environments. Nat Genet 40, 471–475. 10.1038/ng.110

Álvarez-Arenas, A., Podolski-Renic, A., Belmonte-Beitia, J., Pesic, M., Calvo, G.F., 2019. Interplay of Darwinian Selection, Lamarckian Induction and Microvesicle Transfer on Drug Resistance in Cancer. Sci Rep 9, 9332. 10.1038/s41598-019-45863-z

Bhang, H.E.C., Ruddy, D.A., Radhakrishna, V.K., Caushi, J.X., Zhao, R., Hims, M.M., Singh, A.P., Kao, I., Rakiec, D., Shaw, P., Balak, M., Raza, A., Ackley, E., Keen, N., Schlabach, M.R., Palmer, M., Leary, R.J., Chiang, D.Y., Sellers, W.R., Michor, F., Cooke, V.G., Korn, J.M., Stegmeier, F., 2015. Studying clonal dynamics in response to cancer therapy using high-complexity barcoding. Nature Medicine 21, 440–448. 10.1038/nm.3841

Bhattarai-kline, S., Lear, S.K., Fishman, C.B., Lopez, S.C., Elana, R., 2022. Recording gene expression order in DNA by CRISPR addition of retron barcodes. 10.1038/s41586-022-04994-6

Blundell, J.R., Levy, S.F., 2014. Beyond genome sequencing: Lineage tracking with barcodes to study the dynamics of evolution, infection, and cancer. Genomics 104, 1–14. 10.1016/j.ygeno.2014.09.005

Blundell, J.R., Schwartz, K., Francois, D., Fisher, D.S., Sherlock, G., Levy, S.F., 2019. The dynamics of adaptive genetic diversity during the early stages of clonal evolution. Nature Ecology and Evolution 3, 293–301. 10.1038/s41559-018-0758-1

Cheng, F.H.C., Aguda, B.D., Tsai, J.-C., Kochańczyk, M., Lin, J.M.J., Chen, G.C.W., Lai, H.-C., Nephew, K.P., Hwang, T.-W., Chan, M.W.Y., 2014. A Mathematical Model of Bimodal Epigenetic Control of miR-193a in Ovarian Cancer Stem Cells. PLOS ONE 9, e116050. 10.1371/journal.pone.0116050

Chmielecki, J., Foo, J., Oxnard, G.R., Hutchinson, K., Ohashi, K., Somwar, R., Wang, L., Amato, K.R., Arcila, M., Sos, M.L., Socci, N.D., Viale, A., De Stanchina, E., Ginsberg, M.S., Thomas, R.K., Kris, M.G., Inoue, A., Ladanyi, M., Miller, V.A., Michor, F., Pao, W., 2011. Optimization of dosing for EGFR-mutant non-small cell lung cancer with evolutionary cancer modeling. Science Translational Medicine 3. 10.1126/scitranslmed.3002356

Dhawan, A., Madani Tonekaboni, S.A., Taube, J.H., Hu, S., Sphyris, N., Mani, S.A., Kohandel, M., 2016. Mathematical modelling of phenotypic plasticity and conversion to a stem-cell state under hypoxia. Sci Rep 6, 18074. 10.1038/srep18074

Diaz Jr, L.A., Williams, R.T., Wu, J., Kinde, I., Hecht, J.R., Berlin, J., Allen, B., Bozic, I., Reiter, J.G., Nowak, M.A., Kinzler, K.W., Oliner, K.S., Vogelstein, B., 2012. The molecular evolution of acquired resistance to targeted EGFR blockade in colorectal cancers. Nature 486, 537–540. 10.1038/nature11219

Ding, L., Ley, T.J., Larson, D.E., Miller, C.A., Koboldt, D.C., Welch, J.S., Ritchey, J.K., Young, M.A., Lamprecht, T., McLellan, M.D., McMichael, J.F., Wallis, J.W., Lu, C., Shen, D., Harris, C.C., Dooling, D.J., Fulton, R.S., Fulton, L.L., Chen, K., Schmidt, H., Kalicki-Veizer, J., Magrini, V.J., Cook, L., McGrath, S.D., Vickery, T.L., Wendl, M.C., Heath, S., Watson, M.A., Link, D.C., Tomasson, M.H., Shannon, W.D., Payton, J.E., Kulkarni, S., Westervelt, P., Walter, M.J., Graubert, T.A., Mardis, E.R., Wilson, R.K., Dipersio, J.F., 2012. Clonal evolution in relapsed acute myeloid leukaemia revealed by whole-genome sequencing. Nature 481, 506–510. 10.1038/nature10738

Emert, B.L., Cote, C., Torre, E.A., Dardani, I.P., Jiang, C.L., Jain, N., Shaffer, S.M., Raj, A., 2021. Variability within rare cell states enables multiple paths toward drug resistance. Nature Biotechnology 39, 865–876. 10.1038/s41587-021-00837-3

Eyler, C.E., Matsunaga, H., Hovestadt, V., Vantine, S.J., Van Galen, P., Bernstein, B.E., 2020. Single-cell lineage analysis reveals genetic and epigenetic interplay in glioblastoma drug resistance. Genome Biology 21, 1–21. 10.1186/s13059-020-02085-1

Fabre, M.A., de Almeida, J.G., Fiorillo, E., Mitchell, E., Damaskou, A., Rak, J., Orrù, V., Marongiu, M., Chapman, M.S., Vijayabaskar, M.S., Baxter, J., Hardy, C., Abascal, F., Williams, N., Nangalia, J., Martincorena, I., Campbell, P.J., McKinney, E.F., Cucca, F., Gerstung, M., Vassiliou, G.S., 2022. The longitudinal dynamics and natural history of clonal haematopoiesis. Nature. 10.1038/s41586-022-04785-z

Foo, J., Michor, F., 2009. Evolution of resistance to targeted anti-cancer therapies during continuous and pulsed administration strategies. PLoS Computational Biology 5. 10.1371/journal.pcbi.1000557

Francois, O., Laval, G., 2011. Deviance Information Criteria for Model Selection in Approximate Bayesian Computation. Statistical Applications in Genetics and Molecular Biology 10. 10.2202/1544-6115.1678

Gabbutt, C., Schenck, R.O., Weisenberger, D.J., Kimberley, C., Berner, A., Househam, J., Lakatos, E., Robertson-Tessi, M., Martin, I., Patel, R., Clark, S.K., Latchford, A., Barnes, C.P., Leedham, S.J., Anderson, A.R.A., Graham, T.A., Shibata, D., 2022a. Fluctuating methylation clocks for cell lineage tracing at high temporal resolution in human tissues. Nat Biotechnol 40, 720–730. 10.1038/s41587-021-01109-w

Gabbutt, C., Wright, N.A., Baker, A., Shibata, D., Graham, T.A., 2022b. Lineage tracing in human tissues. J Pathol 257, 501–512. 10.1002/path.5911

Gallagher, K., Strobl, M.A., Park, D.S., Spoendlin, F.C., Gatenby, R.A., Maini, P.K., Anderson, A.R., 2024. Mathematical Model-Driven Deep Learning Enables Personalized Adaptive Therapy. Cancer Research. 10.1158/0008-5472.CAN-23-2040

Germano, D.P.J., Zarebski, A.E., Hautphenne, S., Moss, R., Flegg, J.A., Flegg, M.B., 2024. Jump-Switch-Flow: hybrid stochastic-deterministic solutions of compartmental models. 10.48550/arXiv.2405.13239

Gunnarsson, E.B., De, S., Leder, K., Foo, J., 2020. Understanding the role of phenotypic switching in cancer drug resistance. Journal of Theoretical Biology 490, 110162. 10.1016/j.jtbi.2020.110162

Hari, K., Sabuwala, B., Subramani, B.V., La Porta, C.A.M., Zapperi, S., Font-Clos, F., Jolly, M.K., 2020. Identifying inhibitors of epithelial–mesenchymal plasticity using a network topology-based approach. npj Syst Biol Appl 6, 1–12. 10.1038/s41540-020-0132-1

Hinohara, K., Wu, H.J., Vigneau, S., McDonald, T.O., Igarashi, K.J., Yamamoto, K.N., Madsen, T., Fassl, A., Egri, S.B., Papanastasiou, M., Ding, L., Peluffo, G., Cohen, O., Kales, S.C., Lal-Nag, M., Rai, G., Maloney, D.J., Jadhav, A., Simeonov, A., Wagle, N., Brown, M., Meissner, A., Sicinski, P., Jaffe, J.D., Jeselsohn, R., Gimelbrant, A.A., Michor, F., Polyak, K., 2018. KDM5 Histone Demethylase Activity Links Cellular Transcriptomic Heterogeneity to Therapeutic Resistance. Cancer Cell 34, 939–953.e9. 10.1016/j.ccell.2018.10.014

Howard, G.R., Johnson, K.E., Rodriguez Ayala, A., Yankeelov, T.E., Brock, A., 2018. A multi-state model of chemoresistance to characterize phenotypic dynamics in breast cancer. Scientific Reports 8, 1–11. 10.1038/s41598-018-30467-w

Jasinska, W., Manhart, M., Lerner, J., Gauthier, L., Serohijos, A.W.R., Bershtein, S., 2020. Chromosomal barcoding of E. coli populations reveals lineage diversity dynamics at high resolution. Nature Ecology and Evolution 4, 437–452. 10.1038/s41559-020-1103-z

Jensen, N.F., Stenvang, J., Beck, M.K., Hanáková, B., Belling, K.C., Do, K.N., Viuff, B., Nygård, S.B., Gupta, R., Rasmussen, M.H., Tarpgaard, L.S., Hansen, T.P., Budinská, E., Pfeiffer, P., Bosman, F., Tejpar, S., Roth, A., Delorenzi, M., Andersen, C.L., Rømer, M.U., Brünner, N., Moreira, J.M.A., 2015. Establishment and characterization of models of chemotherapy resistance in colorectal cancer: Towards a predictive signature of chemoresistance. Mol Oncol 9, 1169–1185. 10.1016/j.molonc.2015.02.008

Johnson, K.E., Howard, G.R., Morgan, D., Brenner, E.A., Gardner, A.L., Durrett, R.E., Mo, W., Al’khafaji, A., Sontag, E.D., Jarrett, A.M., Yankeelov, T.E., Brock, A., 2020. Integrating transcriptomics and bulk time course data into a mathematical framework to describe and predict therapeutic resistance in cancer. Physical Biology 18. 10.1088/1478-3975/abb09c

Jost, L., 2006. Entropy and diversity. Oikos 113, 363–375.

Kalhor, R., Mali, P., Church, G.M., 2017. Rapidly evolving homing CRISPR barcodes. Nat Methods 14, 195–200. 10.1038/nmeth.4108

Kebschull, J.M., Zador, A.M., 2018. Cellular barcoding: lineage tracing, screening and beyond. Nature Methods 15, 871–879. 10.1038/s41592-018-0185-x

Kreger, J., Komarova, N.L., Wodarz, D., 2021. A hybrid stochastic-deterministic approach to explore multiple infection and evolution in HIV. PLOS Computational Biology 17, e1009713. 10.1371/journal.pcbi.1009713

Kuosmanen, T., Cairns, J., Noble, R., Beerenwinkel, N., Mononen, T., Mustonen, V., 2021. Drug-induced resistance evolution necessitates less aggressive treatment. PLoS Comput Biol 17, e1009418. 10.1371/journal.pcbi.1009418

Lan, X., Jörg, D.J., Cavalli, F.M.G., Richards, L.M., Nguyen, L.V., Vanner, R.J., Guilhamon, P., Lee, L., Kushida, M.M., Pellacani, D., Park, N.I., Coutinho, F.J., Whetstone, H., Selvadurai, H.J., Che, C., Luu, B., Carles, A., Moksa, M., Rastegar, N., Head, R., Dolma, S., Prinos, P., Cusimano, M.D., Das, S., Bernstein, M., Arrowsmith, C.H., Mungall, A.J., Moore, R.A., Ma, Y., Gallo, M., Lupien, M., Pugh, T.J., Taylor, M.D., Hirst, M., Eaves, C.J., Simons, B.D., Dirks, P.B., 2017. Fate mapping of human glioblastoma reveals an invariant stem cell hierarchy. Nature 549, 227–232. 10.1038/nature23666

Levy, S.F., Blundell, J.R., Venkataram, S., Petrov, D.A., Fisher, D.S., Sherlock, G., 2015. Quantitative evolutionary dynamics using high-resolution lineage tracking. Nature 519, 181–186. 10.1038/nature14279

Luria, S.E., Delbrück, M., 1943. Mutations of Bacteria From Virus Sensitivity To Virus Resistance. Genetics 28, 491–511. 10.1093/genetics/28.6.491

Maltas, J., Killarney, S.T., Singleton, K.R., Strobl, M.A.R., Washart, R., Wood, K.C., Wood, K.B., 2024. Drug dependence in cancer is exploitable by optimally constructed treatment holidays. Nat Ecol Evol 8, 147–162. 10.1038/s41559-023-02255-x

Michor, F., Hughes, T.P., Iwasa, Y., Branford, S., Shah, N.P., Sawyers, C.L., Nowak, M.A., 2005. Dynamics of chronic myeloid leukaemia. Nature 435, 1267–1270. 10.1038/nature03669

Misale, S., Bozic, I., Tong, J., Peraza-Penton, A., Lallo, A., Baldi, F., Lin, K.H., Truini, M., Trusolino, L., Bertotti, A., Di Nicolantonio, F., Nowak, M.A., Zhang, L., Wood, K.C., Bardelli, A., 2015. Vertical suppression of the EGFR pathway prevents onset of resistance in colorectal cancers. Nature Communications 6. 10.1038/ncomms9305

Misale, S., Yaeger, R., Hobor, S., Scala, E., Janakiraman, M., Liska, D., Valtorta, E., Schiavo, R., Buscarino, M., Siravegna, G., Bencardino, K., Cercek, A., Chen, C.T., Veronese, S., Zanon, C., Sartore-Bianchi, A., Gambacorta, M., Gallicchio, M., Vakiani, E., Boscaro, V., Medico, E., Weiser, M., Siena, S., Di Nicolantonio, F., Solit, D., Bardelli, A., 2012. Emergence of KRAS mutations and acquired resistance to anti-EGFR therapy in colorectal cancer. Nature 486, 532–536. 10.1038/nature11156

Nam, A.S., Chaligne, R., Landau, D.A., 2021. Integrating genetic and non-genetic determinants of cancer evolution by single-cell multi-omics. Nature Reviews Genetics 22, 3–18. 10.1038/s41576-020-0265-5

Oren, Y., Tsabar, M., Cuoco, M.S., Amir-Zilberstein, L., Cabanos, H.F., Hütter, J.-C., Hu, B., Thakore, P.I., Tabaka, M., Fulco, C.P., Colgan, W., Cuevas, B.M., Hurvitz, S.A., Slamon, D.J., Deik, A., Pierce, K.A., Clish, C., Hata, A.N., Zaganjor, E., Lahav, G., Politi, K., Brugge, J.S., Regev, A., 2021. Cycling cancer persister cells arise from lineages with distinct programs. Nature. 10.1038/s41586-021-03796-6

Pisco, A.O., Brock, A., Zhou, J., Moor, A., Mojtahedi, M., Jackson, D., Huang, S., 2013. Non-Darwinian dynamics in therapy-induced cancer drug resistance. Nature Communications 4, 1–11. 10.1038/ncomms3467

Poels, K.E., Schoenfeld, A.J., Makhnin, A., Tobi, Y., Wang, Y., Frisco-Cabanos, H., Chakrabarti, S., Shi, M., Napoli, C., McDonald, T.O., Tan, W., Hata, A., Weinrich, S.L., Yu, H.A., Michor, F., 2021. Identification of optimal dosing schedules of dacomitinib and osimertinib for a phase I/II trial in advanced EGFR-mutant non-small cell lung cancer. Nat Commun 12, 3697. 10.1038/s41467-021-23912-4

Ramirez, M., Rajaram, S., Steininger, R.J., Osipchuk, D., Roth, M.A., Morinishi, L.S., Evans, L., Ji, W., Hsu, C.H., Thurley, K., Wei, S., Zhou, A., Koduru, P.R., Posner, B.A., Wu, L.F., Altschuler, S.J., 2016. Diverse drug-resistance mechanisms can emerge from drug-tolerant cancer persister cells. Nature Communications 7, 1–8. 10.1038/ncomms10690

Rehman, S.K., Haynes, J., Collignon, E., Brown, K.R., Wang, Y., Nixon, A.M.L., Bruce, J.P., Wintersinger, J.A., Singh Mer, A., Lo, E.B.L., Leung, C., Lima-Fernandes, E., Pedley, N.M., Soares, F., McGibbon, S., He, H.H., Pollet, A., Pugh, T.J., Haibe-Kains, B., Morris, Q., Ramalho-Santos, M., Goyal, S., Moffat, J., O’Brien, C.A., 2021. Colorectal Cancer Cells Enter a Diapause-like DTP State to Survive Chemotherapy. Cell 184, 226–242.e21. 10.1016/j.cell.2020.11.018

Roswell, M., Dushoff, J., Winfree, R., 2020. A conceptual guide to measuring species diversity. Oikos 128, 659–667. 10.1111/oik.07202

Russo, M., Pompei, S., Sogari, A., Corigliano, M., Crisafulli, G., Puliafito, A., Lamba, S., Erriquez, J., Bertotti, A., Gherardi, M., Di Nicolantonio, F., Bardelli, A., Cosentino Lagomarsino, M., 2022. A modified fluctuation-test framework characterizes the population dynamics and mutation rate of colorectal cancer persister cells. Nat Genet 54, 976–984. 10.1038/s41588-022-01105-z

Shaffer, S.M., Dunagin, M.C., Torborg, S.R., Torre, E.A., Emert, B., Krepler, C., Beqiri, M., Sproesser, K., Brafford, P.A., Xiao, M., Eggan, E., Anastopoulos, I.N., Vargas-Garcia, C.A., Singh, A., Nathanson, K.L., Herlyn, M., Raj, A., 2017. Rare cell variability and drug-induced reprogramming as a mode of cancer drug resistance. Nature 546, 431–435. 10.1038/nature22794

Shaffer, S.M., Emert, B.L., Reyes Hueros, R.A., Cote, C., Harmange, G., Schaff, D.L., Sizemore, A.E., Gupte, R., Torre, E., Singh, A., Bassett, D.S., Raj, A., 2020. Memory Sequencing Reveals Heritable Single-Cell Gene Expression Programs Associated with Distinct Cellular Behaviors. Cell 182, 947–959.e17. 10.1016/j.cell.2020.07.003

Sharma, S.V., Lee, D.Y., Li, B., Quinlan, M.P., Takahashi, F., Maheswaran, S., McDermott, U., Azizian, N., Zou, L., Fischbach, M.A., Wong, K.K., Brandstetter, K., Wittner, B., Ramaswamy, S., Classon, M., Settleman, J., 2010. A Chromatin-Mediated Reversible Drug-Tolerant State in Cancer Cell Subpopulations. Cell 141, 69–80. 10.1016/j.cell.2010.02.027

Shi, H., Hugo, W., Kong, X., Hong, A., Koya, R.C., Moriceau, G., Chodon, T., Guo, R., Johnson, D.B., Dahlman, K.B., Kelley, M.C., Kefford, R.F., Chmielowski, B., Glaspy, J.A., Sosman, J.A., Van Baren, N., Long, G.V., Ribas, A., Lo, R.S., 2014. Acquired resistance and clonal evolution in melanoma during BRAF inhibitor therapy. Cancer Discovery 4. 10.1158/2159-8290.CD-13-0642

Skoulidis, F., Li, B.T., Dy, G.K., Price, T.J., Falchook, G.S., Wolf, J., Italiano, A., Schuler, M., Borghaei, H., Barlesi, F., Kato, T., Curioni-Fontecedro, A., Sacher, A., Spira, A., Ramalingam, S.S., Takahashi, T., Besse, B., Anderson, A., Ang, A., Tran, Q., Mather, O., Henary, H., Ngarmchamnanrith, G., Friberg, G., Velcheti, V., Govindan, R., 2021. Sotorasib for Lung Cancers with *KRAS* p.G12C Mutation. N Engl J Med 384, 2371– 2381. 10.1056/NEJMoa2103695

Soltani, M., Vargas-Garcia, C.A., Antunes, D., Singh, A., 2016. Intercellular Variability in Protein Levels from Stochastic Expression and Noisy Cell Cycle Processes. PLOS Computational Biology 12, e1004972. 10.1371/journal.pcbi.1004972

Tadrowski, A.C., Evans, M.R., Waclaw, B., 2018. Phenotypic Switching Can Speed up Microbial Evolution. Scientific Reports 8, 8941. 10.1038/s41598-018-27095-9

Viossat, Y., Noble, R., 2021. A theoretical analysis of tumour containment. Nature Ecology & Evolution. 10.1038/s41559-021-01428-w

Warner, E.W., Van der Eecken, K., Murtha, A.J., Kwan, E.M., Herberts, C., Sipola, J., Ng, S.W.S., Chen, X.E., Fonseca, N.M., Ritch, E., Schönlau, E., Bernales, C.Q., Donnellan, G., Munzur, A.D., Parekh, K., Beja, K., Wong, A., Verbeke, S., Lumen, N., Van Dorpe, J., De Laere, B., Annala, M., Vandekerkhove, G., Ost, P., Wyatt, A.W., 2024. Multiregion sampling of de novo metastatic prostate cancer reveals complex polyclonality and augments clinical genotyping. Nat Cancer 5, 114–130. 10.1038/s43018-023-00692-y

West, J., Ma, Y., Newton, P.K., 2018. Capitalizing on competition: An evolutionary model of competitive release in metastatic castration resistant prostate cancer treatment. Journal of Theoretical Biology 455, 249–260. 10.1016/j.jtbi.2018.07.028

Windels, E.M., Michiels, J.E., Fauvart, M., Wenseleers, T., Van den Bergh, B., Michiels, J., 2019. Bacterial persistence promotes the evolution of antibiotic resistance by increasing survival and mutation rates. ISME Journal 13, 1239–1251. 10.1038/s41396-019-0344-9

Yang, D., Jones, M.G., Naranjo, S., Yosef, N., Jacks, T., Weissman, J.S., 2022. Lineage tracing reveals the phylodynamics, plasticity, and paths of tumor evolution Lineage tracing reveals the phylodynamics, plasticity, and paths of tumor evolution. Cell 1–19. 10.1016/j.cell.2022.04.015

Zahn, H., Steif, A., Laks, E., Eirew, P., VanInsberghe, M., Shah, S.P., Aparicio, S., Hansen, C.L., 2017. Scalable whole-genome single-cell library preparation without preamplification. Nat Methods 14, 167–173. 10.1038/nmeth.4140

## Methods References

Daniel Lai, G.H. (2017) ‘HMMcopy’. Bioconductor. Available at: 10.18129/B9.BIOC.HMMCOPY.

Funnell, T. et al. (2022) ‘Single-cell genomic variation induced by mutational processes in cancer’, Nature, 612(7938), pp. 106–115. Available at: 10.1038/s41586-022-05249-0.

Gaspar, J.M. (2018) ‘NGmerge: merging paired-end reads via novel empirically-derived models of sequencing errors’, BMC Bioinformatics, 19(1), p. 536. Available at: 10.1186/s12859-018-2579-2.

Hao, Y. et al. (2021) ‘Integrated analysis of multimodal single-cell data’, Cell, 184(13), pp. 3573–3587.e29. Available at: 10.1016/j.cell.2021.04.048.

Korotkevich, G. et al. (2016) *Fast gene set enrichment analysis*. preprint. Bioinformatics. Available at: 10.1101/060012.

McGinnis, C.S., Murrow, L.M. and Gartner, Z.J. (2019) ‘DoubletFinder: Doublet Detection in Single-Cell RNA Sequencing Data Using Artificial Nearest Neighbors’, Cell Systems, 8(4), pp. 329–337.e4. Available at: 10.1016/j.cels.2019.03.003.

Muyas, F. et al. (2023) ‘De novo detection of somatic mutations in high-throughput single-cell profiling data sets’, Nature Biotechnology [Preprint]. Available at: 10.1038/s41587-023-01863-z.

Ritchie, M.E. et al. (2015) ‘limma powers differential expression analyses for RNA-sequencing and microarray studies’, Nucleic Acids Research, 43(7), pp. e47–e47. Available at: 10.1093/nar/gkv007.

Robinson, M.D., McCarthy, D.J. and Smyth, G.K. (2010) ‘edgeR : a Bioconductor package for differential expression analysis of digital gene expression data’, Bioinformatics, 26(1), pp. 139–140. Available at: 10.1093/bioinformatics/btp616.

Zhao, L. et al. (2018) ‘Bartender: A fast and accurate clustering algorithm to count barcode reads’, Bioinformatics, 34(5), pp. 739–747. Available at: 10.1093/bioinformatics/btx655.

