## Supplementary Figures for "Quantitative measurement of phenotype dynamics during cancer drug resistance evolution using genetic barcoding"

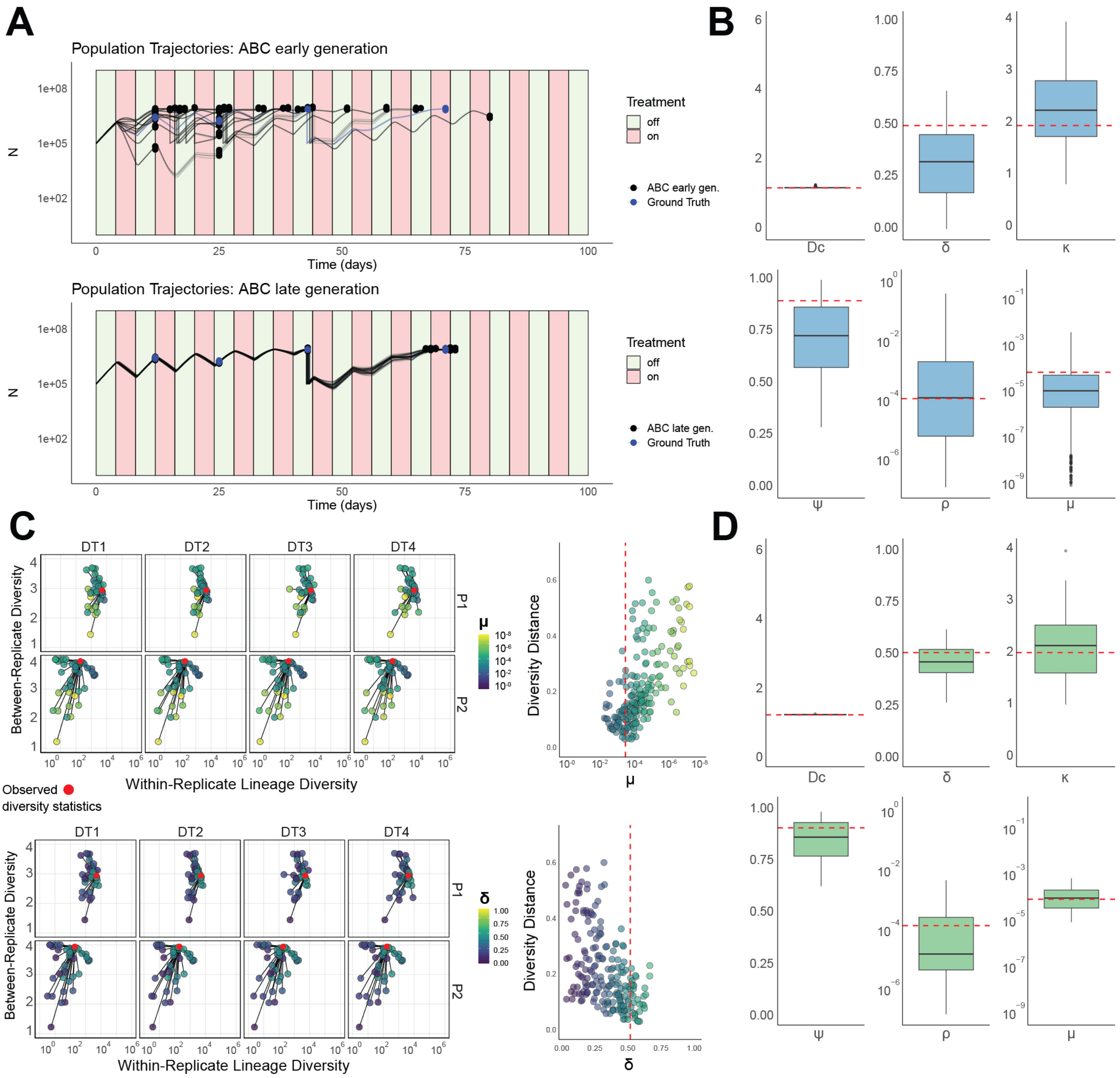
A
Population Trajectories: ABC early generation
Treatment
off
on
ABC early gen.
Ground Truth
Time (days)
N
Population Trajectories: ABC late generation
ABC late gen.
Ground Truth
B
Dc
δ
κ
ψ
ρ
μ
C
DT1
DT2
DT3
DT4
P1
P2
Between-Replicate Diversity
Within-Replicate Lineage Diversity
Observed diversity statistics
Diversity Distance
D

**Supplementary Fig. 1 | Incorporating lineage data alongside population size changes increases the power to recover parameters governing resistance evolution. A)** Population size changes through treatment simulated using posterior predictive parameters from an early (top) and late (bottom) generation of the approximate Bayesian computation (ABC) inference first step that fits the total cell size changes at a set of given times to observed replicate timepoints (points). **B)** Posterior distributions (n=500, 4 experimental replicates) of the inferred parameter values using only the observed population size changes (true parameter values - red dashed lines). **C)** A subset of the simulated lineage diversity statistics for each drug-treatment replicate's (DT1-4) two timepoints (P1-2) used for the parameter inference step, highlighted by one of the model parameters (Top: sensitive to resistant phenotype transition probability per cell-division - $\mu$) and the fitness penalty of resistance in the absence of treatment (Bottom: controlled by $\delta$) compared to the true diversity statistics for the given simulation (red points). Adjacent panels show the lineage diversity distance as a function of the two parameters, with the true value highlighted (red dashed line). **D)** Posterior distributions (n=50, 4 experimental replicates) of the inferred parameter values using the combined cell population size and cell lineage statistics (true parameter values – red dashed lines). Boxplots show the median, the first and third quartiles, and whiskers 1.5x the interquartile range. Parameters: $\rho$ - pre-existing fraction of resistance, $\mu$ – sensitive to resistant transition probability per cell division, $\psi$ – strength of the resistant phenotype, Dc – maximum strength of the drug, $\kappa$ – accumulation/decay rate of the drug.

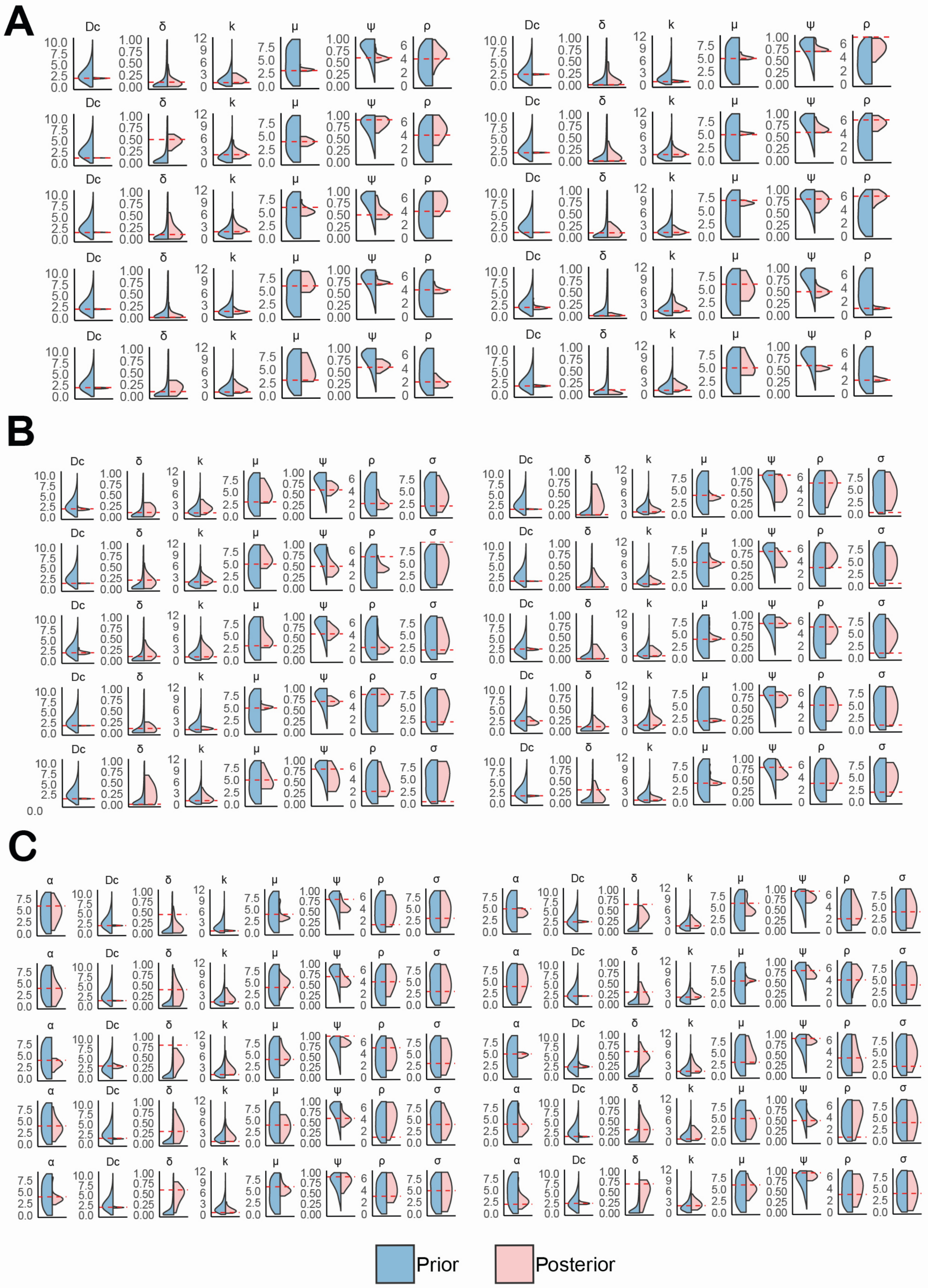
A
B
C
Dc
δ
k
μ
ψ
ρ
σ
α
Prior
Posterior

**Supplementary Fig. 2 | Simulated posterior distributions given different resistance models and evolutionary scenarios.** Prior vs posterior distributions given a representative set of simulated synthetic data using our full inference framework (both population trajectories and lineage distributions) for all three phenotype evolution models considered: **A)** uni-directional transitions (Model A), **B)** bi-directional transitions (Model B), **C)** escape transitions (Model C). Parameters: $\rho$ - pre-existing fraction of resistance, $\mu$ – sensitive to resistant transition probability per cell division, $\sigma$ – resistant to sensitive transition probability per cell division, $\alpha$ – resistant to escape transition probability per cell division, $\psi$ – strength of the resistant phenotype, Dc – maximum strength of the drug, $\kappa$ – accumulation/decay rate of the drug. The following parameters are shown as -log10($x$) : $\rho$, $\mu$, $\sigma$, $\alpha$.

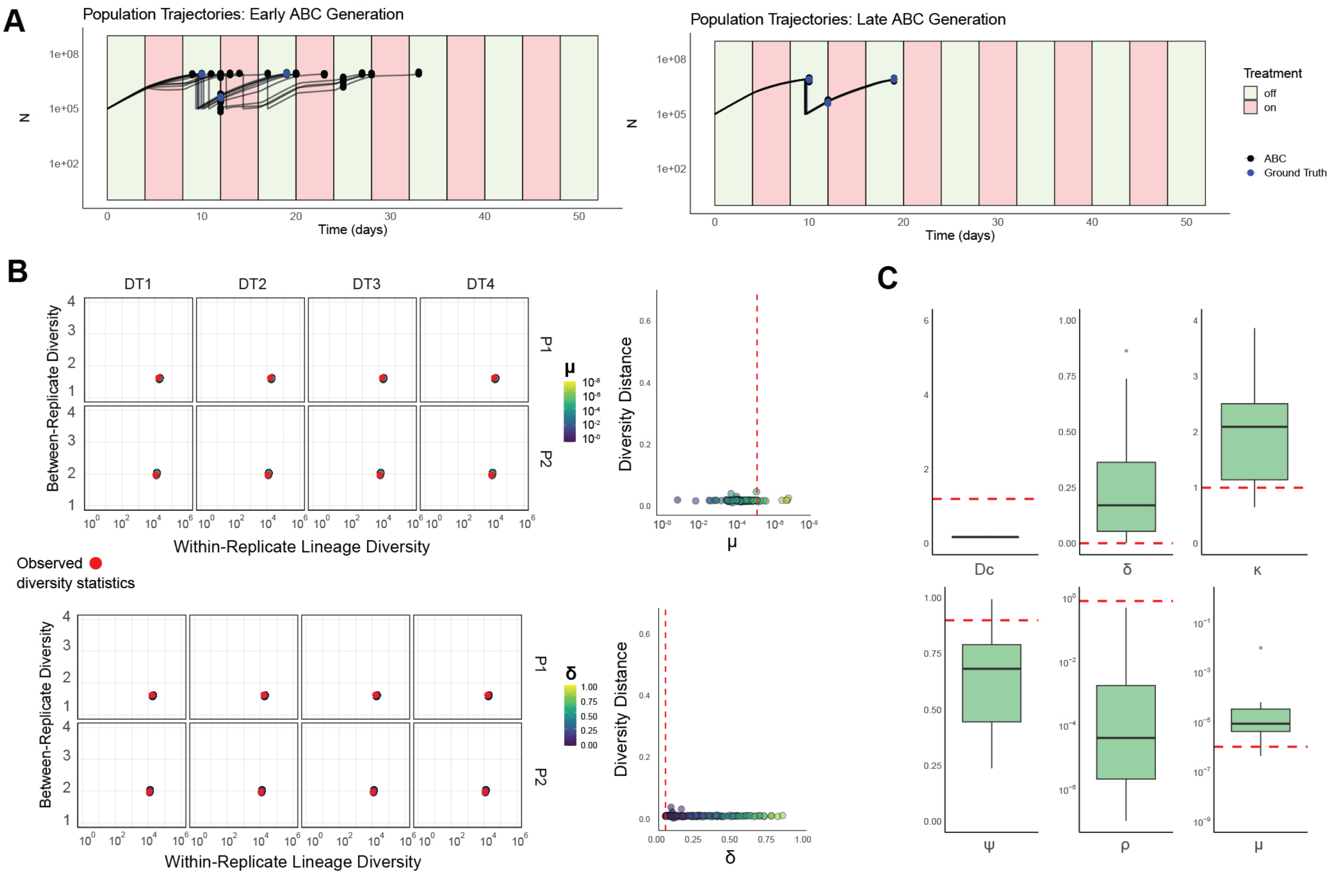
A
Population Trajectories: Early ABC Generation
Population Trajectories: Late ABC Generation
N
1e+08
1e+05
1e+02
Time (days)
Treatment
off
on
ABC
Ground Truth
B
DT1
DT2
DT3
DT4
P1
P2
Between-Replicate Diversity
Within-Replicate Lineage Diversity
Observed diversity statistics
μ
δ
Diversity Distance
C
Dc
δ
κ
ψ
ρ
μ

**Supplementary Fig. 3 | The modelling framework struggles to recover the true parameter values when resistance is very common and/or the effect of treatment is weak. A)** Population size changes through treatment simulated using posterior predictive parameters from an early (left) and late (right) generation of the approximate Bayesian computation (ABC) inference first step that fits the total cell size changes at a set of given times to observed replicate timepoints (points). **B)** A subset of the simulated lineage diversity statistics for each replicate's (DT1-4) two timepoints (P1-2) used for the parameter inference step, highlighted by one of the model parameters (Top: sensitive to resistant phenotype transition probability per cell-division – $\mu$. Bottom: the fitness penalty of resistance in the absence of treatment - $\delta$) compared to the true diversity statistics for the given simulation (red points). Adjacent panels show the lineage diversity distance as a function of the two parameters, with the true value highlighted (red dashed line). **C)** Posterior distributions (n=50, 4 experimental replicates) of the inferred parameter values using the combined cell population size and cell lineage statistics (true parameter values – red dashed lines). Boxplots show the median, the first and third quartiles, and whiskers 1.5x the interquartile range. Parameters: $\rho$ - pre-existing fraction of resistance, $\mu$ – sensitive to resistant transition probability per cell division, $\psi$ – strength of the resistant phenotype, Dc – maximum strength of the drug, $\kappa$ – accumulation/decay rate of the drug.

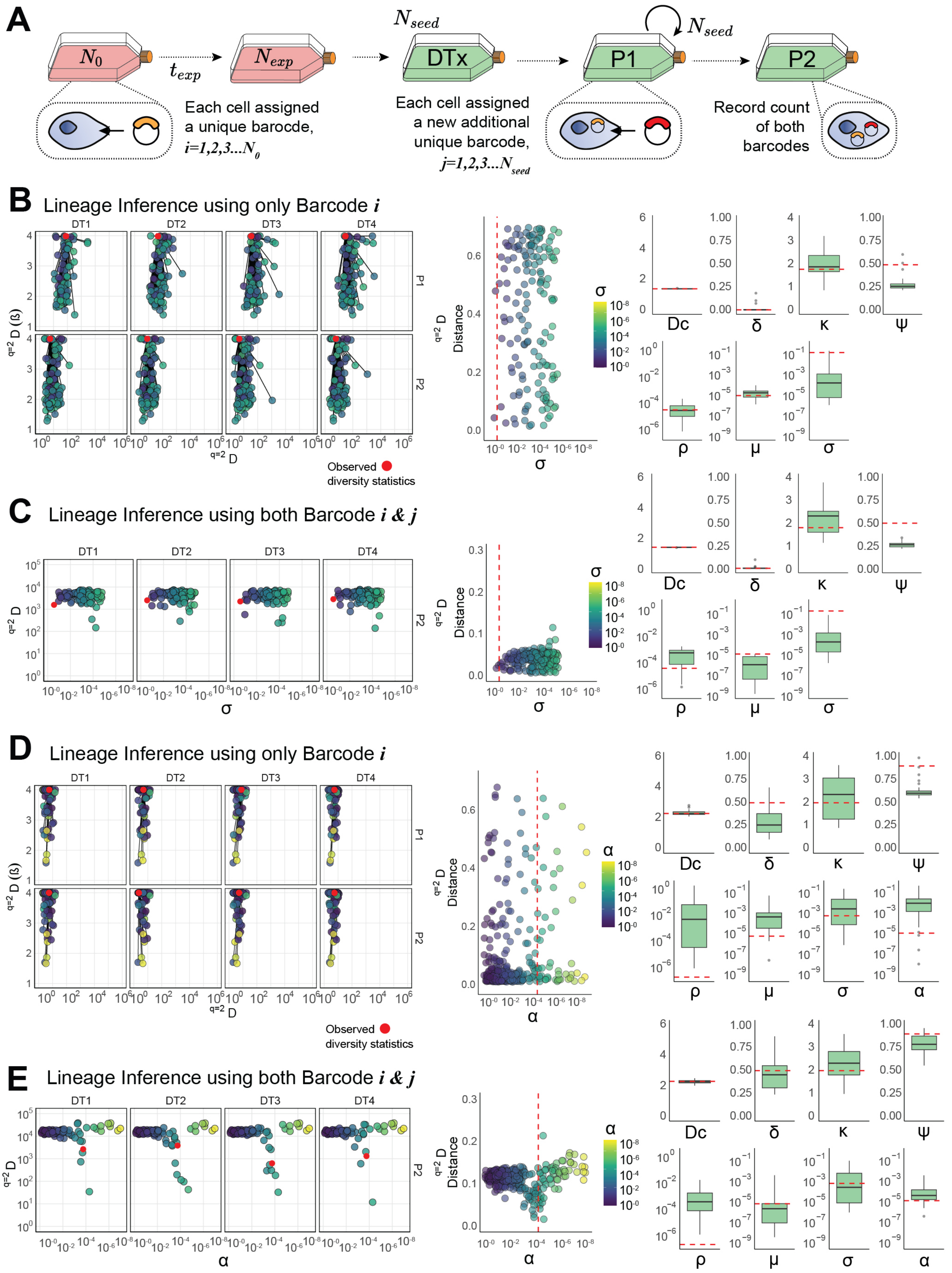
A
$N_0$
$t_{exp}$
$N_{exp}$
$N_{seed}$
DTx
P1
$N_{seed}$
P2
Each cell assigned a unique barocde, i=1,2,3...$N_0$
Each cell assigned a new additional unique barcode, j=1,2,3...$N_{seed}$
Record count of both barcodes
B
Lineage Inference using only Barcode i
DT1
DT2
DT3
DT4
P1
P2
q=2 D (ß)
q=2 D
Observed diversity statistics
Distance
σ
Dc
δ
κ
Ψ
ρ
μ
C
Lineage Inference using both Barcode i & j
D
Lineage Inference using only Barcode i
α
E
Lineage Inference using both Barcode i & j

**Supplementary Fig. 4 | A single re-barcoding step can improve the recovery of true parameter values under certain conditions. A)** A schematic illustrating how cells were barcoded with a second lineage barcode tag at the time of P1. The counts of both barcodes were recorded at the end of the simulation. Within- and between-replicate diversity was calculated for barcode *i*, whereas only within-replicate diversity was calculated for barcode *j*. **B)** The lineage diversity statistics for each replicate's (DT1-4) Passage (P1-2) for a simulation using Model B (bidirectional transitions) calculated using the first barcode comparing simulated to observed values as a function of the reversion transition parameter ($\sigma$). The distances in diversity space are summarised in the middle column and the posterior estimates (n=50, 4 experimental replicates) for all Model B parameters are shown on the RHS with red dashed lines indicating the true values. **C)** As in (B), but now only showing the within-replicate diversity (qD) calculated using the second barcode (*j*). **D)** As in (B) but a simulation using Model C (escape transitions). **E)** As in (C) but a simulation using Model C (escape transitions). Boxplots show the median, the first and third quartiles, and whiskers 1.5x the interquartile range. Parameters: $\rho$ - pre-existing fraction of resistance, $\mu$ – sensitive to resistant transition probability per cell division, $\sigma$ – resistant to sensitive transition probability per cell division, $\alpha$ – resistant to escape transition probability per cell division, $\psi$ – strength of the resistant phenotype, Dc – maximum strength of the drug, $\kappa$ – accumulation/decay rate of the drug.

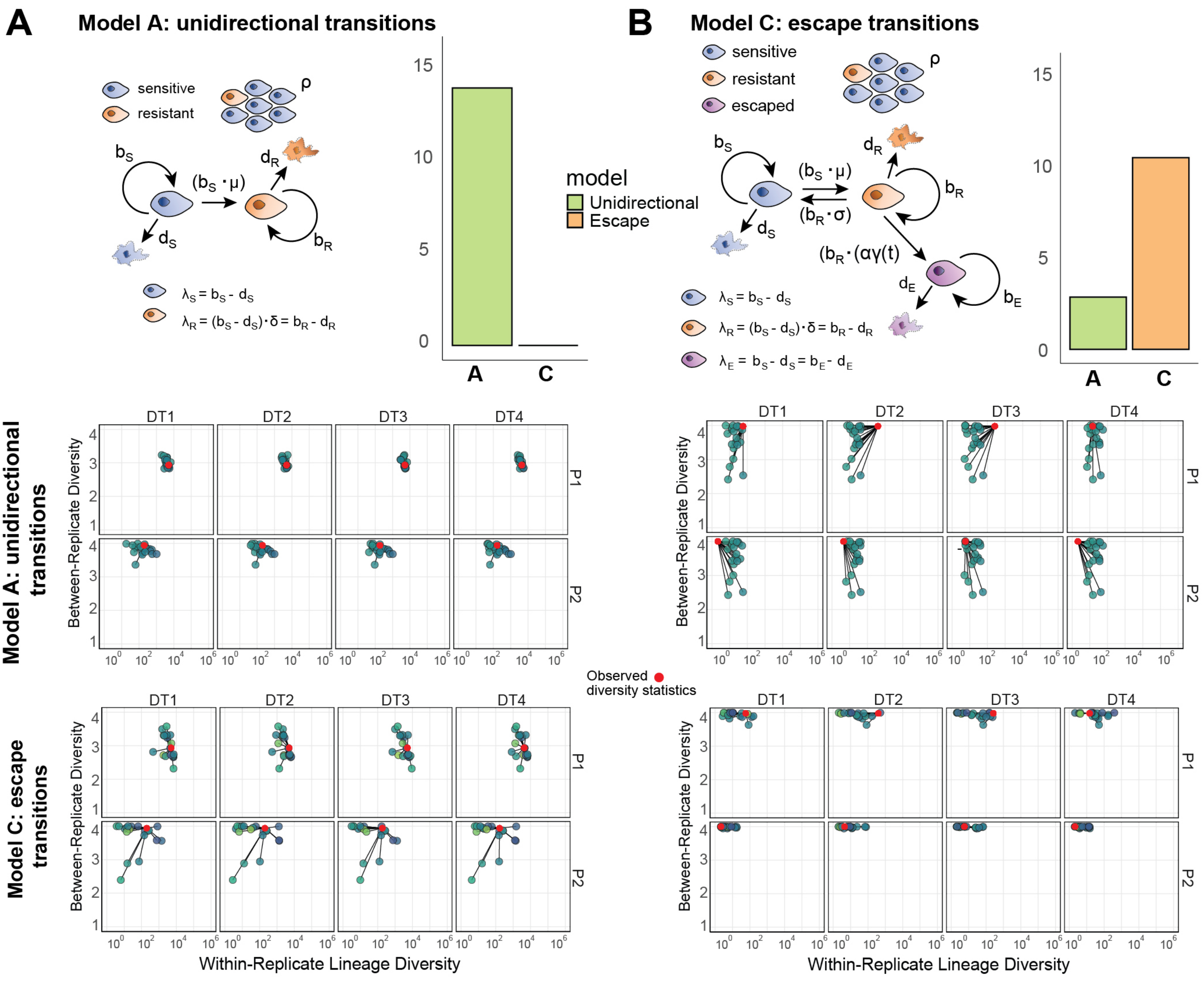
A
Model A: unidirectional transitions
sensitive
resistant
ρ
model
Unidirectional
Escape
B
Model C: escape transitions
sensitive
resistant
escaped
Model A: unidirectional transitions
Model C: escape transitions
Between-Replicate Diversity
Within-Replicate Lineage Diversity
DT1
DT2
DT3
DT4
P1
P2
Observed diversity statistics

**Supplementary Fig. 5 | Model selection with DIC scores.** The number of times the model selection step chose either of the two models, given the true underlying model was Model A, unidirectional transitions (**A**) or Model C, escape transitions (**B**). Schematics and parameter values are shown for our first (unidirectional switching, LHS) and third (escape transition, RHS) models (**A**&**B**, respectively): the columns corresponds to the model that generated the data for the results shown, whereas the rows correspond to the model fit to the data. A subset of diversity statistics are shown for posterior predictive simulations of each model for the unidirectional model (Model A - top panels) and escape model (Model C - bottom panels). For each of the example outputs shown the lower DIC score supports the correct model.

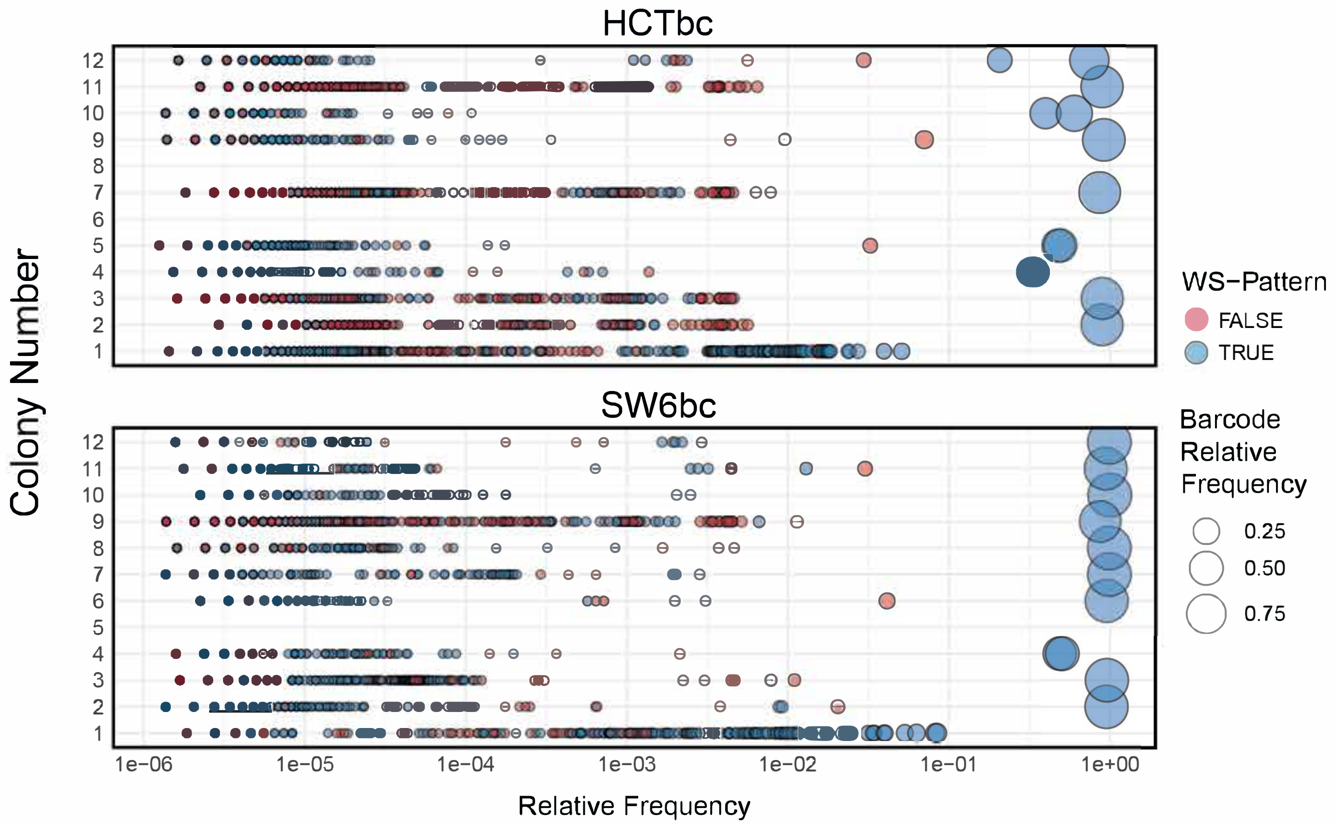
HCTbc
SW6bc
Colony Number
Relative Frequency
1e−06
1e−05
1e−04
1e−03
1e−02
1e−01
1e+00
WS−Pattern
FALSE
TRUE
Barcode
Relative
Frequency
0.25
0.50
0.75

**Supplementary Fig. 6 | The majority of cells contain a single barcode which adheres to the expected nucleotide pattern.** The relative frequency of sequenced barcodes in expanded single cell colonies from each of our barcoded cell lines (top: HCTbc, bottom: SW6bc) highlighted by whether or not the sequenced barcode adhered to the expected ClonTracer weak-strong 30bp nucleotide pattern. Colony 1 corresponds to a control well where 100 cells were seeded and expanded before sequencing.

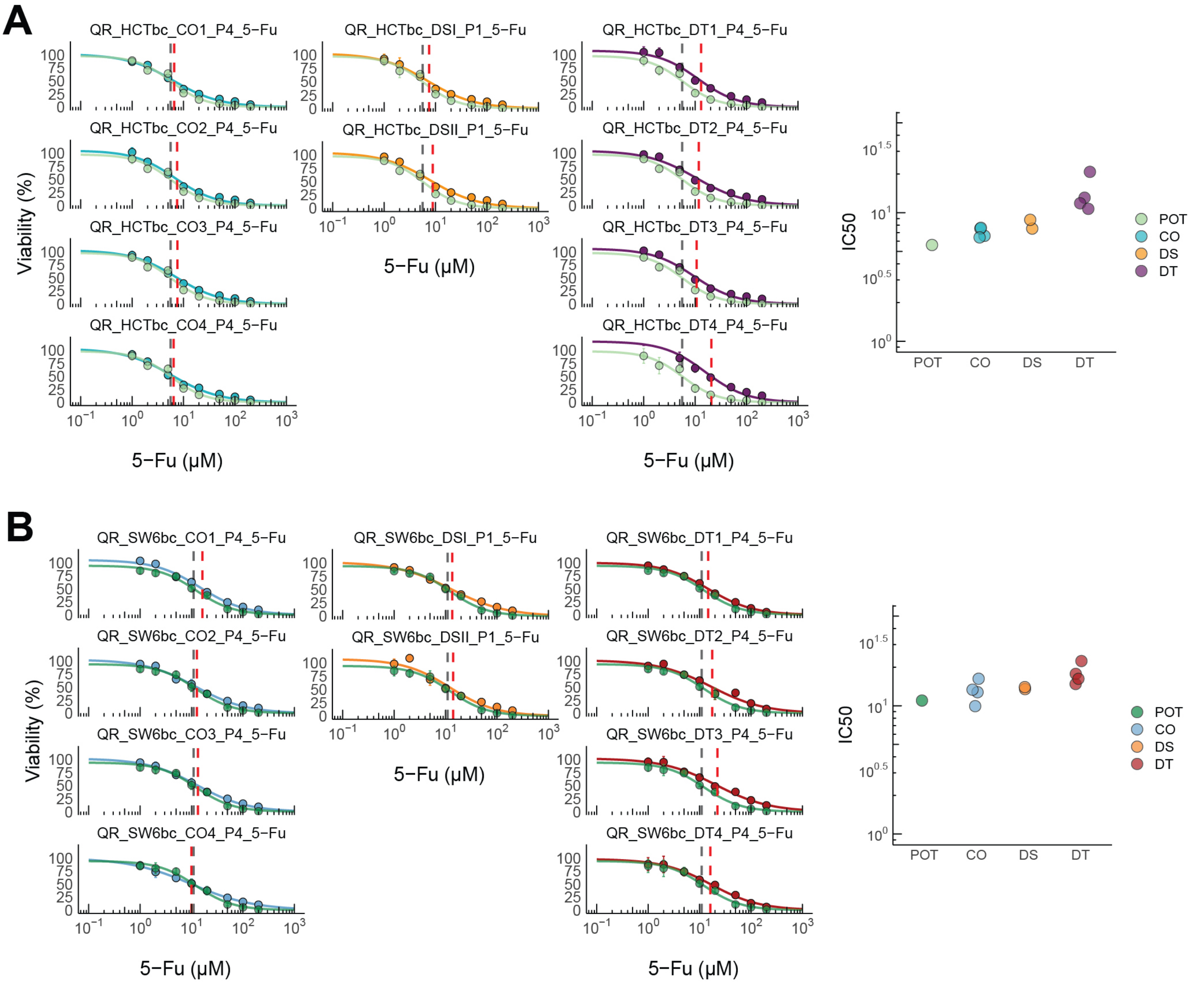
A
QR_HCTbc_CO1_P4_5−Fu
QR_HCTbc_CO2_P4_5−Fu
QR_HCTbc_CO3_P4_5−Fu
QR_HCTbc_CO4_P4_5−Fu
QR_HCTbc_DSI_P1_5−Fu
QR_HCTbc_DSII_P1_5−Fu
QR_HCTbc_DT1_P4_5−Fu
QR_HCTbc_DT2_P4_5−Fu
QR_HCTbc_DT3_P4_5−Fu
QR_HCTbc_DT4_P4_5−Fu
Viability (%)
5−Fu (μM)
IC50
POT
CO
DS
DT
B
QR_SW6bc_CO1_P4_5−Fu
QR_SW6bc_CO2_P4_5−Fu
QR_SW6bc_CO3_P4_5−Fu
QR_SW6bc_CO4_P4_5−Fu
QR_SW6bc_DSI_P1_5−Fu
QR_SW6bc_DSII_P1_5−Fu
QR_SW6bc_DT1_P4_5−Fu
QR_SW6bc_DT2_P4_5−Fu
QR_SW6bc_DT3_P4_5−Fu
QR_SW6bc_DT4_P4_5−Fu
Viability (%)
5−Fu (μM)
IC50
POT
CO
DS
DT

**Supplementary Fig. 7 | Long-term resistance evolution experiment dose-response curves.** Dose-response curves for all experimental conditions from the P4 samples for each of our barcoded cell lines (top: HCTbc, bottom: SW6bc). Each panel shows the response for the parental (POT) and a given experimental condition (CO: control, DS: drug-stop, DT: drug treatment). Adjacent panels show the IC50 values for all conditions.

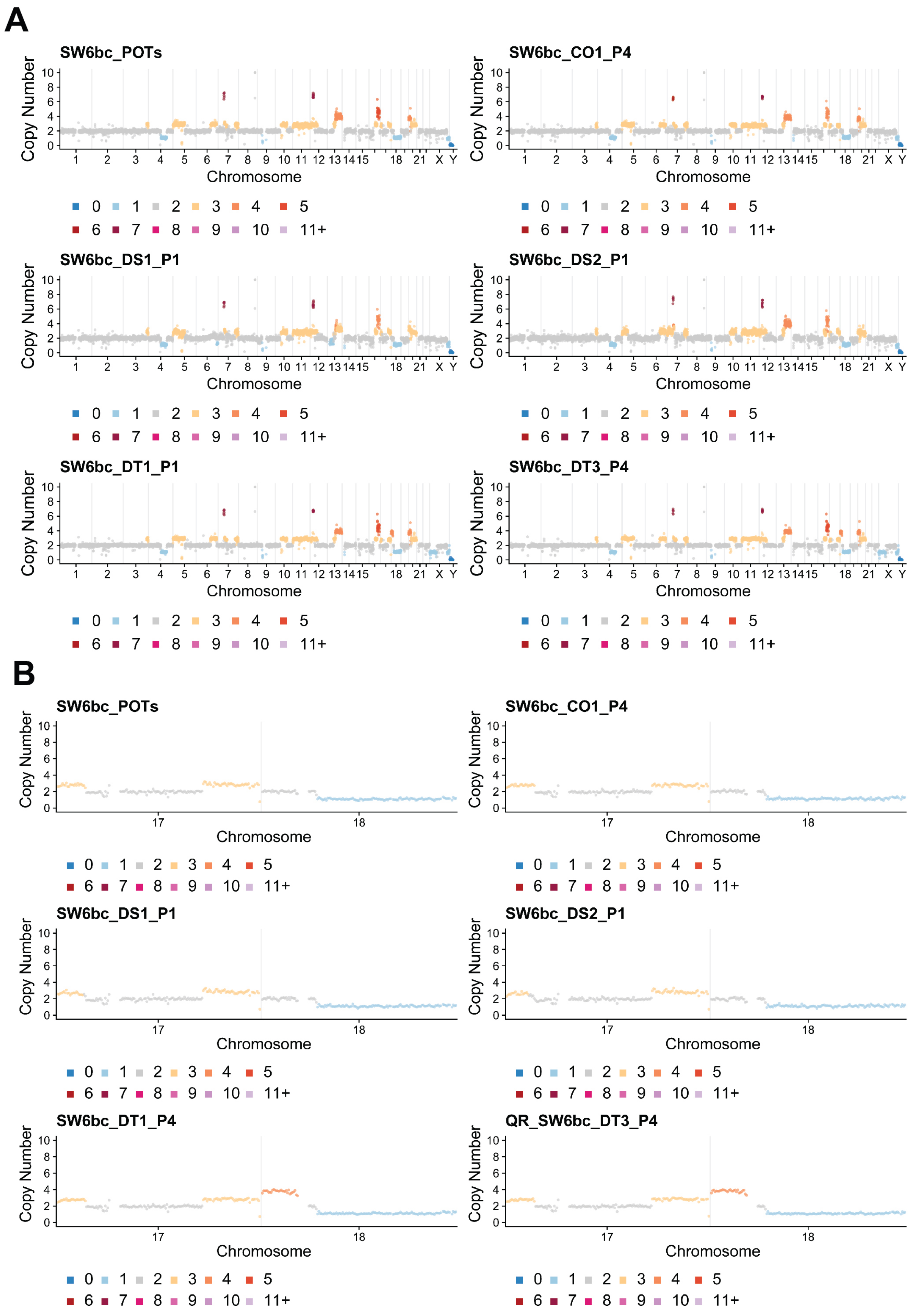
A
SW6bc_POTs
SW6bc_CO1_P4
SW6bc_DS1_P1
SW6bc_DS2_P1
SW6bc_DT1_P1
SW6bc_DT3_P4
Copy Number
0 2 4 6 8 10
1 2 3 4 5 6 7 8 9 10 11 12 13 14 15 18 21 X Y
Chromosome
0 1 2 3 4 5
6 7 8 9 10 11+
B
SW6bc_POTs
SW6bc_CO1_P4
SW6bc_DS1_P1
SW6bc_DS2_P1
SW6bc_DT1_P4
QR_SW6bc_DT3_P4
Copy Number
0 2 4 6 8 10
17 18
Chromosome
0 1 2 3 4 5
6 7 8 9 10 11+

**Supplementary Fig. 8 | Consensus copy number profiles from pooled scWGS of SW6bc experiment replicates. A)** Consensus copy numbers for all chromosomes for the six condition replicates. Each panel shows the consensus copy number for a given experimental condition (CO: control, DS: drug-stop, DT: drug treatment). **B)** Regions on chromosome 17 and 18 for the same condition replicates.

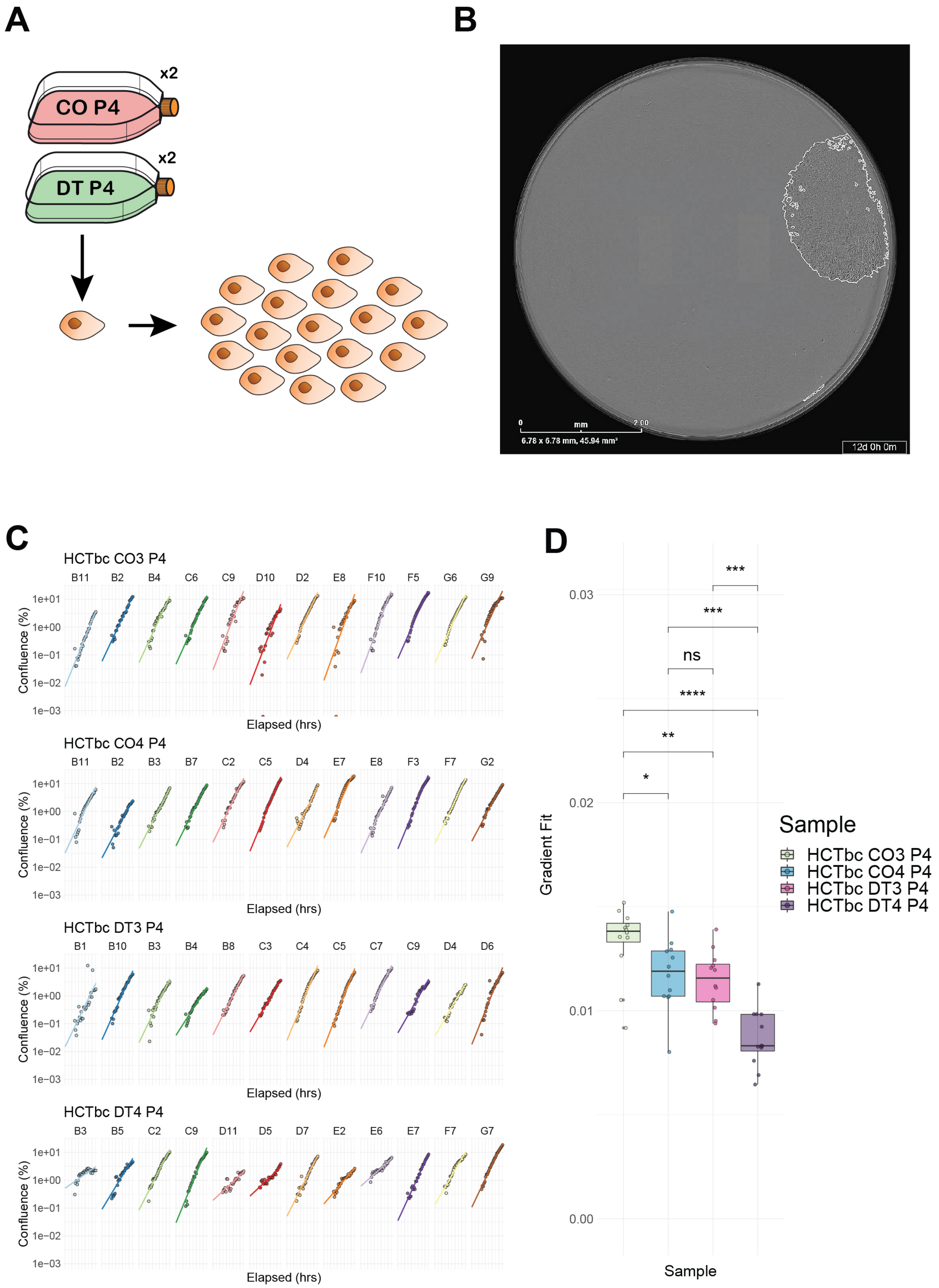
A
B
x2
CO P4
x2
DT P4
0
mm
2.00
6.78 x 6.78 mm, 45.94 mm²
12d 0h 0m
C
HCTbc CO3 P4
B11 B2 B4 C6 C9 D10 D2 E8 F10 F5 G6 G9
Confluence (%)
Elapsed (hrs)
HCTbc CO4 P4
B11 B2 B3 B7 C2 C5 D4 E7 E8 F3 F7 G2
HCTbc DT3 P4
B1 B10 B3 B4 B8 C3 C4 C5 C7 C9 D4 D6
HCTbc DT4 P4
B3 B5 C2 C9 D11 D5 D7 E2 E6 E7 F7 G7
1e+01
1e+00
1e-01
1e-02
1e-03
D
Gradient Fit
0.03
0.02
0.01
0.00
***
***
ns
****
**
*
Sample
Sample
HCTbc CO3 P4
HCTbc CO4 P4
HCTbc DT3 P4
HCTbc DT4 P4

**Supplementary Fig. 9 | Single cell fitness assays in HCTbc. A)** Single cell isolation and clonal expansion was performed for two control (CO) and two drug-treatment (DT) replicates for the HCTbc cell line. **B)** A snapshot of the Incucyte data used to generate the growth rates from confluence readings over time. **C)** Log-confluence estimates and linear regression fits (coloured lines) for 12 chosen colonies per replicate. **D)** Growth rates as estimated by the gradient fits of the log-confluence values per replicate sample. Pairwise comparisons were conducted using the Wilxocon Rank Sum test. Significance levels are annotated as follows: ns $p > 0.05$, * $p<=0.05$, ** $p<=0.01$, *** $p<=0.001$, **** $p <= 0.0001$.

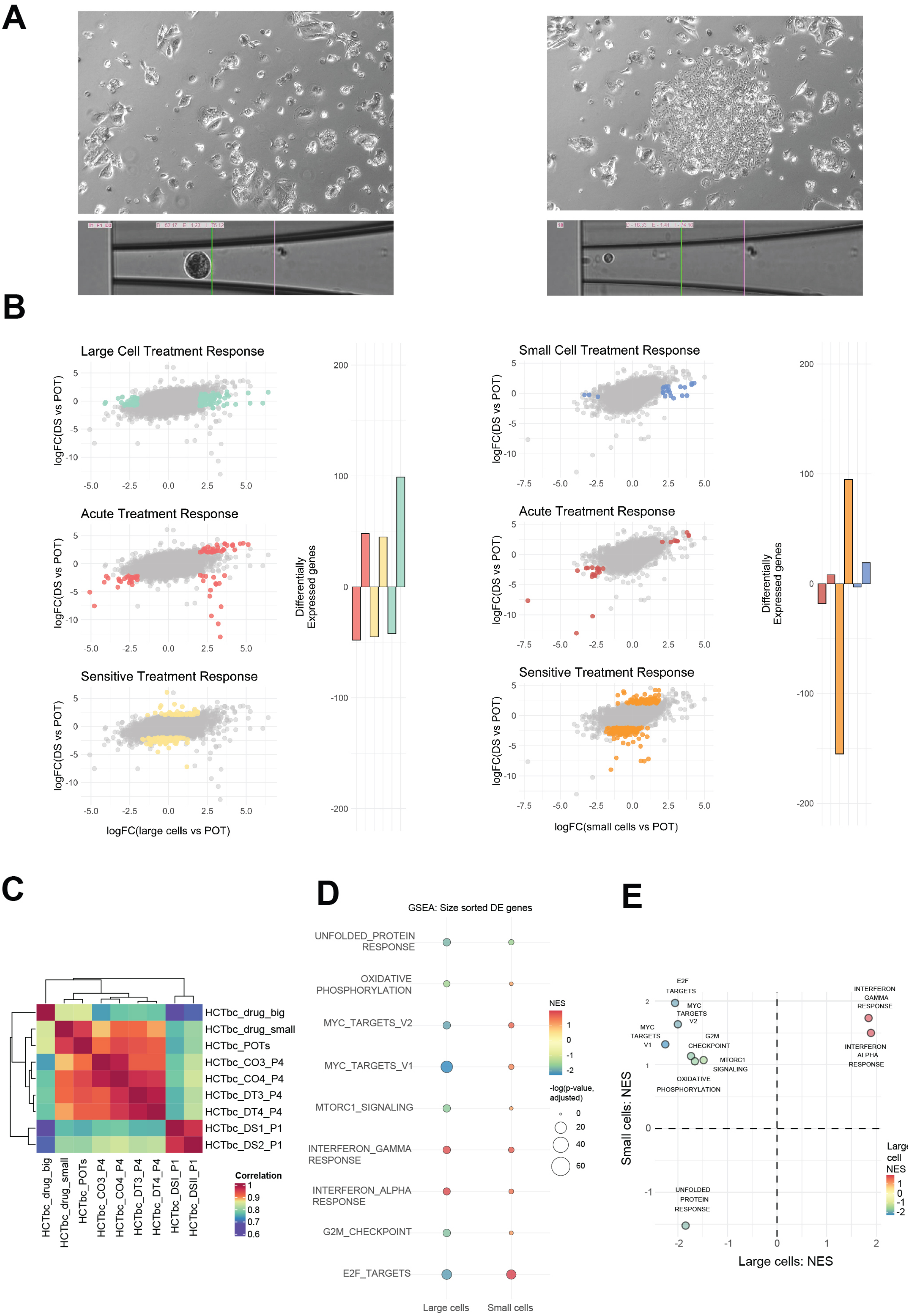
A
B
Large Cell Treatment Response
Acute Treatment Response
Sensitive Treatment Response
logFC(DS vs POT)
logFC(large cells vs POT)
Differentially Expressed genes
Small Cell Treatment Response
Acute Treatment Response
Sensitive Treatment Response
logFC(small cells vs POT)
C
HCTbc_drug_big
HCTbc_drug_small
HCTbc_POTs
HCTbc_CO3_P4
HCTbc_CO4_P4
HCTbc_DT3_P4
HCTbc_DT4_P4
HCTbc_DS1_P1
HCTbc_DS2_P1
Correlation
D
GSEA: Size sorted DE genes
UNFOLDED_PROTEIN RESPONSE
OXIDATIVE PHOSPHORYLATION
MYC_TARGETS_V2
MYC_TARGETS_V1
MTORC1_SIGNALING
INTERFERON_GAMMA RESPONSE
INTERFERON_ALPHA RESPONSE
G2M_CHECKPOINT
E2F_TARGETS
NES
-log(p-value, adjusted)
Large cells
Small cells
E
E2F TARGETS
MYC TARGETS V2
MYC TARGETS V1
G2M CHECKPOINT
MTORC1 SIGNALING
OXIDATIVE PHOSPHORYLATION
INTERFERON GAMMA RESPONSE
INTERFERON ALPHA RESPONSE
UNFOLDED PROTEIN RESPONSE
Small cells: NES
Large cells: NES
Large cell NES

**Supplementary Fig. 10 | Sorting by size reveals distinct transcriptional types in HCTbc cells that survive multiple rounds of treatment. A)** Comparison brightfield images (top panels) of the large (left hand panels) and small (right hand panels) barcoded HCTbc cell phenotypes at the time of single-cell sorting following seven weeks of periodic chemotherapy (5-Fu) treatment with illustrative single cell sorting images (bottom panels). **B)** Genes associated with replicates specific differential expression (DE) identified via specific comparisons (coloured points per panel): Large/SmallTreatment Response - DE in Large/Small cells, but not DS; Acute Treatment Response - DE in Large/Small cells and DS; Sensitive Treatment Response - DE in DS but not Large/Small cells. Comparisons are repeated for each size-sorted phenotype (Large - left panel, Small - right panel) and the number of significantly DE genes in each group are also shown per comparison. Significantly differentially expressed genes were identified using a quasi-likelihood F-test (QLF test) in edgeR. **C)** Correlation coefficients between each replicate given the average expression across all genes and all cells. **D)** Gene set enrichment analysis (GSEA) for each size-sorted phenotype (Large - left column, Small - right column). Only gene sets that were found to be significant (adjusted p-value $< 0.05$) in one of the two phenotypes are shown. Point colour denotes normalised enrichment score (NES). Adjusted p-values were computed using the multilevel adaptive algorithm implemented in fgseaMultilevel. **E)** Difference in NES from the genes GSEA in (D).
